# Evolution-guided engineering of small-molecule biosensors

**DOI:** 10.1101/601823

**Authors:** Tim Snoek, Evan K. Chaberski, Francesca Ambri, Stefan Kol, Sara P. Bjørn, Bo Pang, Jesus F. Barajas, Ditte H. Welner, Michael K. Jensen, Jay D. Keasling

**Author notes:** To whom correspondence should be addressed. Michael K. Jensen.

## Abstract

Allosteric transcription factors (aTFs) have proven widely applicable for biotechnology and synthetic biology as ligand-specific biosensors enabling real-time monitoring, selection and regulation of cellular metabolism. However, both the biosensor specificity and the correlation between ligand concentration and biosensor output signal, also known as the transfer function, often needs to be optimized before meeting application needs. Here, we present a versatile and high-throughput method to evolve and functionalize prokaryotic aTF specificity and transfer functions in a eukaryote chassis, namely baker’s yeast *Saccharomyces cerevisiae*. From a single round of directed evolution of the effector-binding domain (EBD) coupled with various toggled selection regimes, we robustly select aTF variants of the *cis, cis*-muconic acid-inducible transcription factor BenM evolved for change in ligand specificity, increased dynamic output range, shifts in operational range, and a complete inversion of function from activation to repression. Importantly, by targeting only the EBD, the evolved biosensors display DNA-binding affinities similar to BenM, and are functional when ported back into a non-native prokaryote chassis. The developed platform technology thus leverages aTF evolvability for the development of new host-agnostic biosensors with user-defined small-molecule specificities and transfer functions.

## Introduction

The ability to selectively control gene expression has fueled synthetic biology as an engineering discipline. Ever since the description of genetic systems inducible by small molecules, such as IPTG, arabinose or tetracycline^1^, the repertoire of genetic switches for ligand-induced control of gene expression has vastly expanded, targeting diverse applications, including directed evolution of bio-based microbial production, *in situ* diagnosis of human gut microbiota, conditional control of mammalian cell differentiation, and synthetic cell-cell communication devices^2–5^.

Given the large number of allosteric transcription factors (aTFs) present in the prokaryotic kingdom^6^, the diversity of chemical structures recognized^7^, and their modular domain structure encoded by a conserved DNA-binding domain (DBD) linked to a diversified effector-binding domain (EBD)^8^, small-molecule biosensors based on aTFs are a particularly valuable class of genetic switches. Ongoing biosensor research therefore seeks to prospect new biosensors from genomic resources^9, 10^, while also developing general design rules and engineering strategies from existing aTFs. Indeed, due to the modular structure of aTFs, several studies have successfully adopted EBD-swapping strategies into platform DBDs to rationally engineer new aTF logic^11–13^, while engineering EBD destabilization for ligand-controlled biosensor stability also have proven successful^14–16^. However, these rational design strategies for engineering new biosensors suffer from the introduction of cross-talk between ligand specificities, difficulty in creating chimeras from different aTF superfamilies, and the risk of losing allostery^2, 11, 17^. Ultimately, this may impact several aspects of biosensor performance, such as the operational and dynamic output ranges, the specificity, and mode-of-action - collectively referred to as the biosensor transfer function or logic.

Acknowledging that allosteric regulation relies on complex interdomain interactions, studies beyond pure rational engineering have successfully adopted directed evolution based on global and randomised mutagenesis approaches for engineering new aTF logic. For instance, the dynamic range of aTFs, i.e. the quantitative relationship between small-molecule inducer concentration and biosensor output signal, has been optimized by directed evolution to match biosensor performance to the experimental design and application needs^18^. Likewise, when no available biosensor exists to the ligand of interest, attempts on both randomised and structure-guided directed evolution of new biosensor specificities and improved operational ranges from existing aTFs have also been successfully demonstrated^19–22^. Moreover, starting from allosterically-dead aTF variants with constitutive DNA-binding, semi-structure-guided mutagenesis has been used to identify and evolve new biosensors with changes in both dynamic output range and inversion of function (ie. inverse-repression)^23–25^. In most of these library studies, the mutagenized aTF libraries were characterized based on ligand-induced expression of fluorescent reporter genes or antibiotic resistance genes for multiplexed selection of aTF variants responding to the ‘trait’ of interest using biosensor readouts based on fluorescence-activated cell sorting (FACS) or cell survival, respectively.

Even though transfer functions and specificities can be optimized using directed evolution and high-throughput read-outs, mutagenizing allosteric proteins is known to cause abundant loss-of-function mutants related specifically to residues involved in ligand binding and those required for maintaining aTF structures^26, 27^. Likewise, when mutating aTFs, trade-offs between transfer function parameters are frequently observed, such as variants with increased dynamic output ranges are diluted by aTF variants in a constitutive allosterically active state^24^. For this reason several studies have further adopted toggled selection regimes with both positive (ON) and negative (OFF) selection regimes to robustly evolve aTFs with user-defined changes in small-molecule specificity^27, 28^, changes in dynamic and operational ranges^29^, and inversion of function^30^. Such toggled selection regimes can successfully limit, or completely abolish, aTF variants with unintended trade-offs in transfer functions or ligand cross-reactivity. Also, as biosensor library sizes are typically limited by the transformation efficiency, the power of directed evolution enables identification of beneficial aTF mutations to accumulate over iterative rounds of screening from even relatively small library sizes^31^.

Still, no single strategy has demonstrated the evolvability of multiple transfer function parameters and specificity from one existing aTF. Likewise, for all the studies on engineering prokaryotic aTFs as biosensors based on directed evolution, there is no evidence available for the portability of engineered aTFs as functional biosensors between different host organisms^19, 22–24, 27, 30, 32^. From both an engineering and an application point of view, the demonstration of sequence-identifiers related to defined transfer function parameters and specificity, and aTF portability between different hosts is of broad interest for biotechnology and human health applications^7^.

Here, we present a simple and high-throughput method to generate tailor-made aTF biosensors with novel transfer function parameters and specificities by means of a single round of directed evolution coupled to stringent FACS-based toggled selection regimes. Selected archetypical aTF variants display complete affinity-maturation towards a non-cognate ligand, 15-fold increased dynamic output range, a 40-fold shift in operational range, and inversion of function from ligand-induced activation to repression, compared to parental aTF. Furthermore, we sequenced and further characterized selected aTF variants by kinetic analyses and functional studies in order to pinpoint mutations linked to changes in functionality, thereby enabling the identification of mutational hotspots for defined transfer function parameters. The presented method leverages aTF evolvability, and supports the high-throughput development of new biosensors with user-defined small-molecule specificities and transfer functions. Finally, the demonstration that evolved aTF variants can be ported back into bacteria as functional biosensors underscores the general usability of the method for biosensor development for a wide range of hosts.

## Results

### Experimental design and biosensor variant library construction

In order to establish a method for the development of new genetically-encoded biosensors based on allosterically regulated transcription factors (aTFs) with user-defined functionalities, we deployed directed evolution coupled with toggled selection using fluorescence-activated cell sorting (FACS)(Fig. 1A). As for transfer function parameters, we focused on evolving biosensors with quantitative changes in dynamic and operational ranges, qualitative changes related to inversion of function, as well as change of small-molecule specificity. These are frequently targeted parameters in biosensor optimization (Fig. 1B). Furthermore we chose to demonstrate the method in yeast because i) the number of biosensors implemented in yeast, and other eukaryotes, is small as compared to the plethora of ligand-sensing systems available in prokaryotes^7, 33^, and ii) using yeast allowed us to leverage high efficiency of homologous recombination for library generation through plasmid gap repair^18, 34^.

**Fig. 1.**
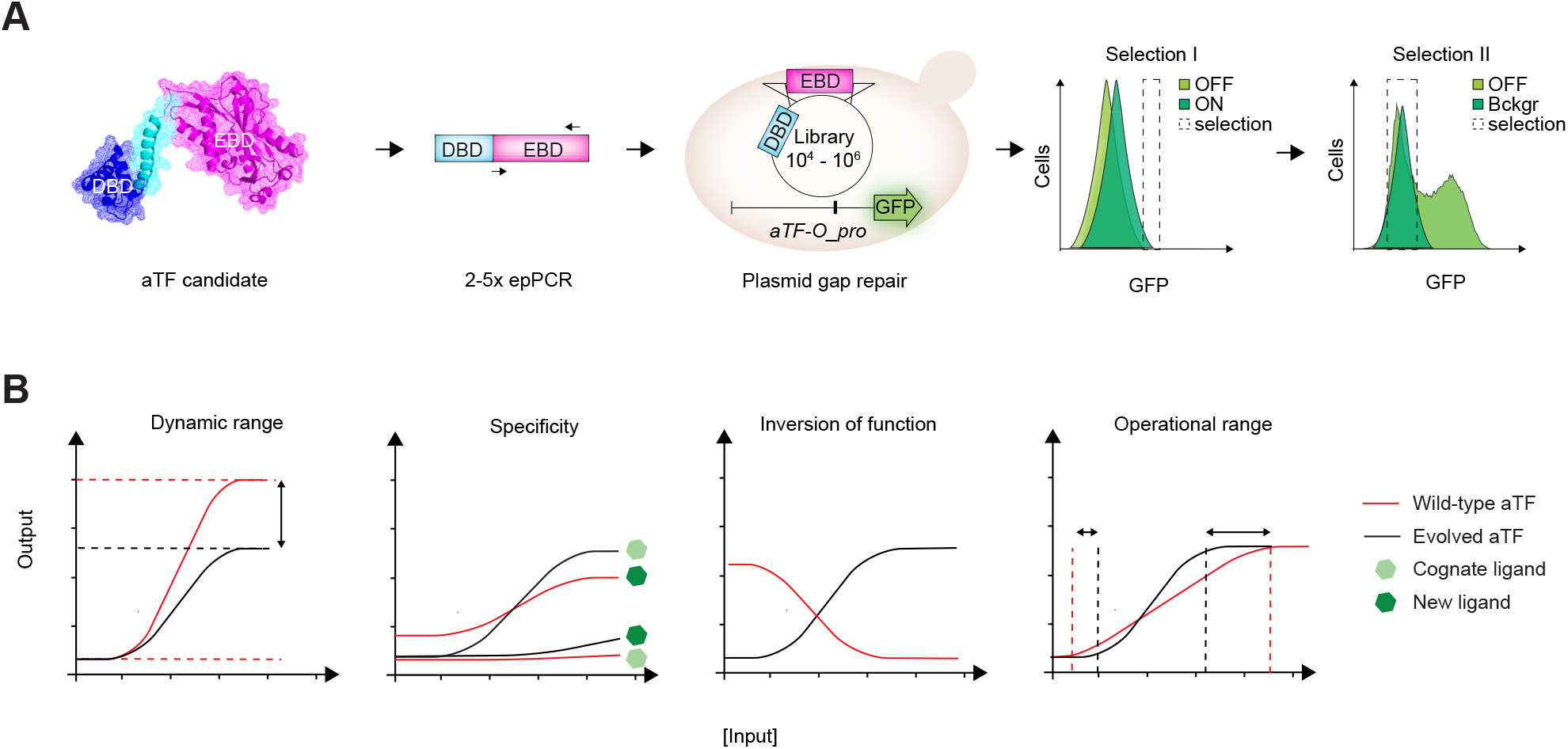
Schematic outline of directed evolution of aTF transfer functions and specificities. (**A**) Overview of experimental workflow using a single round of directed evolution followed by *in vivo* gap reair of aTF variant library in yeast, and finally toggled FACS-based selection of desired transfer function parameters. Protein structure depicts monomeric BenM (PDB 3k1n). (**B**) Illustration of a wild-type aTF transfer function (black) and evolved transfer functions (red) for the traits of interest, namely operational range, dynamic range, and inversion of function, as well as ligand-specificity.

As a proof-of-concept, we used a randomised variant library of the LysR-type *Acinetobacter* sp. ADP1 *cis,cis*-muconic (CCM) acid-binding transcriptional activator BenM, previously engineered as biosensor in the budding yeast *Saccharomyces cerevisiae*^18^. For the aTF variant library generation we considered several aspects in order to generate a high-quality aTF mutant library useful for selection of variants with both changes in transfer function parameters and specificity. First, in order to limit the loss of allostery due to interdomain interactions and to bias the selection towards aTF variants with intact DNA-binding specificity maintained, we specifically focused on evolving the sequence-diverse aTF effector-binding domain (EBD)(Fig. 1A). Next, since we aimed to uncover variants with either quantitative (operational and dynamic ranges) or qualitative (specificity and inversion of function) changes in functionality, we hypothesized a library spanning a variation in mutational loads would be most useful, and therefore created the aTF library consisting of approx 85,000 variants generated from amplicon pools of 2-5 rounds of error-prone PCR (epPCR) targeting the non-conserved EBD residues 73-305 of BenM (Fig. 1A, Suppl. Fig. S1)(see Methods).

For the evolution set-up, the diversified EDB templates were co-transformed with a linearized plasmid backbone encoding the BenM WT DBD, into a platform yeast strain expressing GFP from an engineered weak *CYC1* promoter harboring a previously described aTF binding site (*aTF-O* pro)^18^ to allow for library construction by gap repair and aTF-controlled inducible expression of GFP, respectively (Fig. 1A). For all designs, we genomically integrated *aTF-O::GFP* as a single reporter copy. Next, this library was subjected to various user-defined FACS-based toggled selection regimes in order to evolve aTF variants with new ligand specificity, extended operational and dynamic ranges, and inversion of function (Fig 1A-B). In general first- and second-round sorting included 0.0032 - 5.4% and 34.9 - 43.4% of the library, respectively, depending on the trait sought for (Suppl. Table S1).

Finally, variants evolved in this study were sequenced (including promoter and terminator) and subsequently independently re-transformed into clean yeast background strains, before being re-phenotyped to rule out phenotypic effects of potential secondary genomic mutations.

### Evolution-guided engineering of biosensor transfer function and specificity

The first transfer function parameter to be improved was dynamic range. To this end, we first sorted BenM variants that specifically showed high fluorescent levels in the presence of 1.4 mM CCM (ON state)(Fig. 1B, Fig. 2A, and Suppl. Table S1), a concentration we previously reported relevant for screening in yeast cells^18^. Next, we performed a second round of sorting to remove variants with high background fluorescence in the absence of inducer (OFF state) by sorting variants that did not display background fluorescence under uninduced conditions. Indeed, comparing mean fluorescence intensities of clonal variants isolated after one or two rounds of sorting showed that our toggled selection yielded 62 variants with both lower OFF state (mean OFF state decreased from 4.0 +/- 3.1 to 1.3 +/- 0.6) and higher fold induction (mean fold-induction increased from 4.4 +/- 3.1 to 9.0 +/- 6.1) upon ligand administration as compared to a single round of positive selection (Fig. 2A). More specifically, by sorting 0.17% and 43.4% of our library in ON and OFF states, respectively, we found that 83/94 (88%) of clonal variants obtained after toggled selection improved induction at 1.4 mM CCM with a maximum of 38-fold induction, compared to 2.8-fold induction of BenM WT (Fig. 2A).

**Fig. 2.**
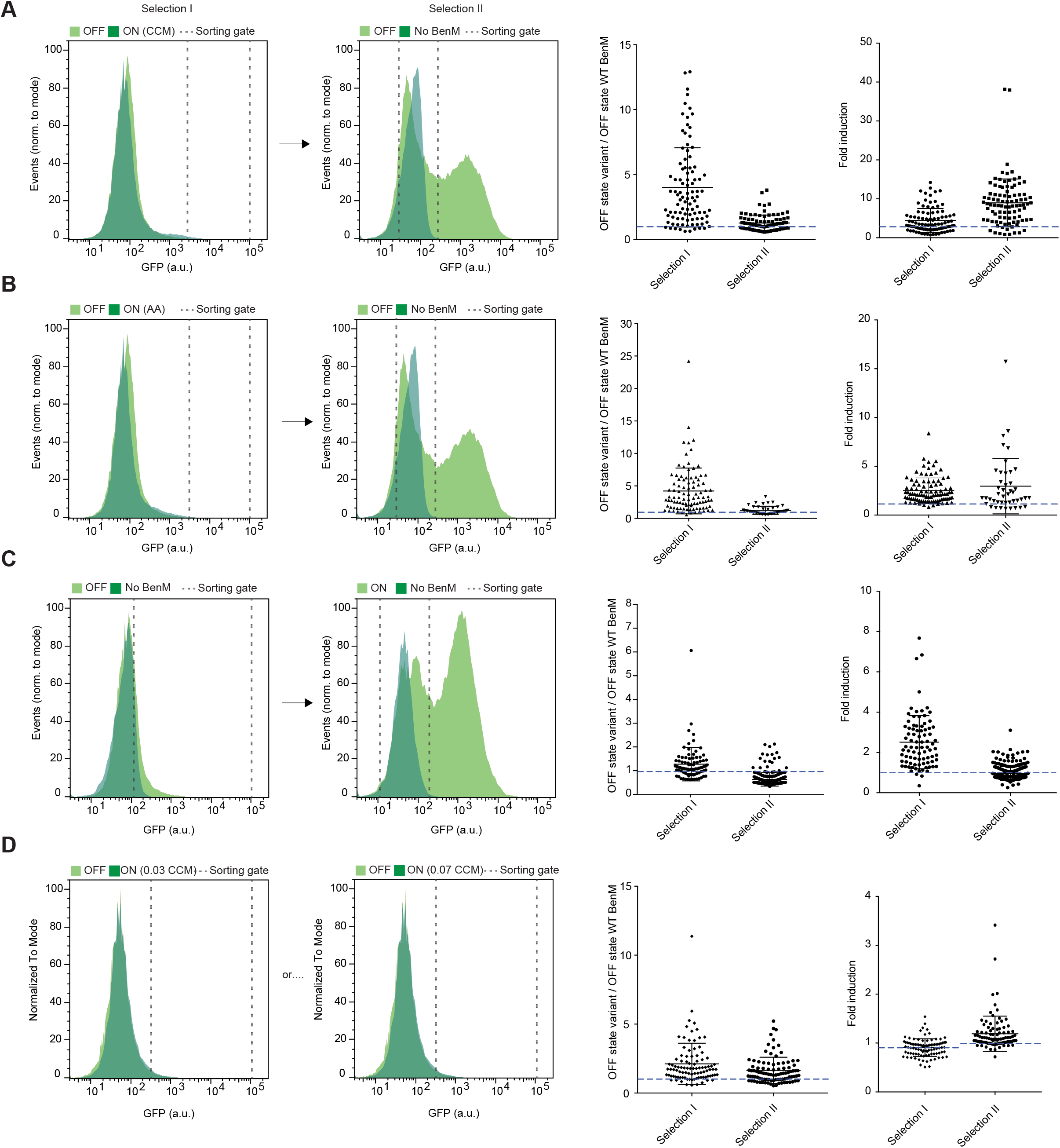
Directed evolution of aTF transfer functions and specificities. *Leftmost and middle-left panels:* Representative flow cytometry green fluorescence protein (GFP) intensity distributions showing the yeast library, or sublibraries thereof, expressing BenM-EBD variants in control (OFF) or inducer medium (ON) and/or a strain not expressing BenM in control medium (No BenM), and dashed lines indicating the sorting gate. (**A**-**B**) *Leftmost and middle-left panels*: Toggled selection for dynamic range (A) or affinity maturation (B). In the first round of selection cells were sorted from the yeast library induced with 1.4 mM CCM (A) or 14 mM adipic acid (B) showing higher fluorescence than the uninduced library. In the second round of selection only variants from the uninduced population were sorted showing background fluorescence, based on a strain not expressing BenM in control medium. *Middle-right panels*: Toggled selection of variants with increased dynamic range in response to CCM (A) and adipic acid (B) show reduced mean OFF state compared to a single round of ON selection. Dashed blue indicate BenM WT OFF state. *Right panels*: Toggled selection of variants with increased dynamic range in response to CCM (A) or change in ligand-specificity (B) towards adipic acid show increased mean fold-induction compared to a single round of ON selection. Dashed blue indicate BenM WT ON state for CCM (2.8x) or adipic acid (1.1x). (**C**) *Leftmost and middle-left panel*: Toggled selection for inversion of function. In the first round of selection cells were sorted from the uninduced yeast library showing higher fluorescence than a strain not expressing BenM in control medium. In the second round of selection only variants from the population induced with 5.6 mM CCM were sorted showing higher than background fluorescence, based on a strain not expressing BenM in control medium. *Middle-right panel*: Toggled selection of clonal variants with inversion of function phenotype show reduced mean OFF state compared to a single round of ON selection. Dashed blue indicate BenM WT OFF state. *Right panel*: Toggled selection of variants with inversion of function phenotype in response to CCM show decreased mean fold-induction compared to a single round of ON selection. Dashed blue indicate a fold-induction of 1.0. (**D**) *Leftmost panels*: Selection for operational range. Cells were sorted from the yeast library induced with 0.035 mM CCM (Selection I) or 0.070 mM CCM (Selection II) showing higher fluorescence than the uninduced library. *Middle-right panel*: Clonal variants gated from inducer media with changed operational range show lowered mean OFF state compared to a single round of positive selection for variants in (A-B). Dashed blue indicate BenM WT OFF state. *Right panel*: Fold-change of variants with changed operational range in response to either 0,035 or 0,07 mM CCM. Dashed blue line indicates fold-induction of WT BenM at the two concentrations (0.90 +/- 0.05 and 1.02 +/- 0.14 for 0.035 mM CCM and 0.070 mM CCM, respectively).

Next, we sought to use directed evolution and toggled selection for affinity-maturation of BenM for new ligand specificity towards adipic acid, a hydrogenated and chemically divergent ligand commercially used in chemical, food and pharmaceutical industries^35^. When engineering new aTF biosensor specificity, it is important to acknowledge that relaxation of ligand specificity towards cognate ligands can impose a challenge in maintaining allostery in transcriptional regulators^16^, and for this reason engineering specificity requires both negative selection (ie. loss of specificity for the native ligand) and positive selection (ie. gain of specificity for the new ligand)^28, 36^. Hence, similar to evolving dynamic range variants, we carried out a toggled selection procedure using adipic acid as an inducer in the ON state, and subsequently sorting variants without background fluorescence under uninduced conditions (OFF state)(Fig. 2B). As we observed when we analyzed clonal variants for changes in dynamic range, we found that the toggled selection procedure reduced the OFF state from a mean value of 4.2 +/- 3.5 to 1.2 +/- 0.6 compared to a single round of positive selection (Fig. 2B). For the affinity-matured variants we found that by sorting 0.032% and 40.5% of our library in ON and OFF states, respectively, a total of 36/44 (82%) clonal variants obtained after toggled selection showed induction by 14 mM adipic acid, with a maximum of 15.7-fold induction (Fig. 2B).

Next, qualitative differences in aTF regulatory mode-of-action suggest that inversion-of-function has occur in evolutionary history. For instance, PurR and TrpR act as repressors when in their ligand bound form^37, 38^, LacI and TetR as repressors only in their apo-form^39, 40^, while BenM acts as a co-activator when bound to CCM^41^. Indeed, previous directed evolution studies have demonstrated the identification of aTFs with reversed mode-of-action^23–25, 30^. To probe the evolvability of BenM towards an inversion-of-function (i.e., deactivator/CCM as a co-repressor) using toggled selection following one round of directed evolution, we first sorted the aTF variant library for highly fluorescent variants in the absence of any ligand, indicative of auto-induction (OFF)(Fig. 2C). Next, we investigated 85 sorted clonal variants by flow cytometry in the presence and absence of inducer. From this screening we found three variants showing decreased reporter gene induction in the presence of CCM (ON). Because of the low success rate of 3.5% (3/85), we tested whether an additional round of sorting for cells that show low fluorescence in the presence of inducer would increase this fraction. From this second screen, we found that 185/285 (65%) after an additional round of negative selection in the presence of 5.6 mM CCM showed a reduction in fluorescence, with one variant showing a 3.9-fold reduction in fluorescence. Overall, mean OFF state for first and second selection regimes were 1.3 +/- 0.7 and 0.6 +/- 0.3, respectively, while mean fold induction was 2.5 +/- 1.3 and 0.9 +/- 0.4, respectively, with the bulk of the second selection variants indeed being deactivated in the presence of CCM (Fig. 2C).

Lastly, a key parameter of biosensor performance is the operational range, i.e. the inducer concentration range in which the sensor shows a significant change in dose-responsive output. Since we found wild-type BenM to show significant induction by CCM from 0.56 mM CCM and upwards, we initially induced the BenM library with 0.035 or 0.070 mM CCM (Fig. 2D). Here, even though the uninduced and induced populations overlapped, we found that a gate based on the top-1% most fluorescent cells under induced conditions (ON) contained, when projected on the uninduced library (OFF), a smaller fraction of the population (i.e., 15% less for 0.035 mM CCM and 20% less for 0.070 mM CCM)(Fig. 2D, Suppl. Table S1), suggesting the presence of variants induced by these low levels of CCM. Subsequent flow cytometry screening of sorted clonal variants showed that for both concentrations the mean OFF state was much lower (2.1 +/- 1.5 for 0.035 mM CCM and 1.6 +/- 1.0 for 0.07 mM CCM (Fig. 2D) than after a single round of positive selection for dynamic range and specificity variants (Fig. 2A-B). Therefore, a second round of sorting to remove auto-inducing variants was deemed unnecessary when evolving operational range variants. In addition, 16/95 (17%) variants for 0.035 mM and 55/95 (58%) variants for 0.070 mM CCM showed a notable fold-induction in the presence of ligand at the concentration they were sorted for (Fig. 2D).

In conclusion, FACS-driven selection regimes for all four parameters yielded new biosensor candidates for desired phenotypes. Toggled selection for dynamic range, affinity-maturation and inversion of function enabled a lowered mean OFF state compared to a single round of selection, whereas the operational range sorting strategy, by design, yielded low-OFF-state variants after a single round of selection. Similarly, fold induction in ON states were higher following toggled selection compared to a single round of selection for dynamic and operational range variants, as well as for the affinity-matured BenM variants. Finally, for inversion-of-function variants toggled selection enabled identification of high OFF state variants with CCM-dependent transcriptional repression.

### Transfer functions and mutation landscapes of evolved aTF variants

To further characterize the evolved variants, sequencing and dose-response experiments of selected isoclonal BenM variants identified from the toggled selection regimes (Fig. 2A-D) were carried out.

By design, our pooled epPCR approach targeted 233 amino acids of BenM, which constitutes the 14-aa DBD-EDB linker, and the two EBD subdomains (I & II) held together by hinge-like β-strands resembling a periplasmic binding protein between which the ligand binds^42^. For the sequence-function analysis, we sequenced 20 BenM mutants, of which three affinity-matured variants were identical. Sequence analysis of the 18 unique BenM variants covering the span of archetypical phenotypes (Fig. 2A-D) identified a total of 75 mutations across 58 positions in the 233-aa window, with each BenM variant having between one (MP17_F12) and six (DAP1_H01) mutations (Fig. 3A-B) illustrative of the pooled epPCR approach undertaken. Moreover, one position (P201S) was mutated in all four specificity variants, two positions (T288I/S and Q291H) were mutated in three variants each, seven positions mutated in two variants each, and the remaining 48 positions mutated in only a single variant (Fig. 3A).

**Fig. 3.**
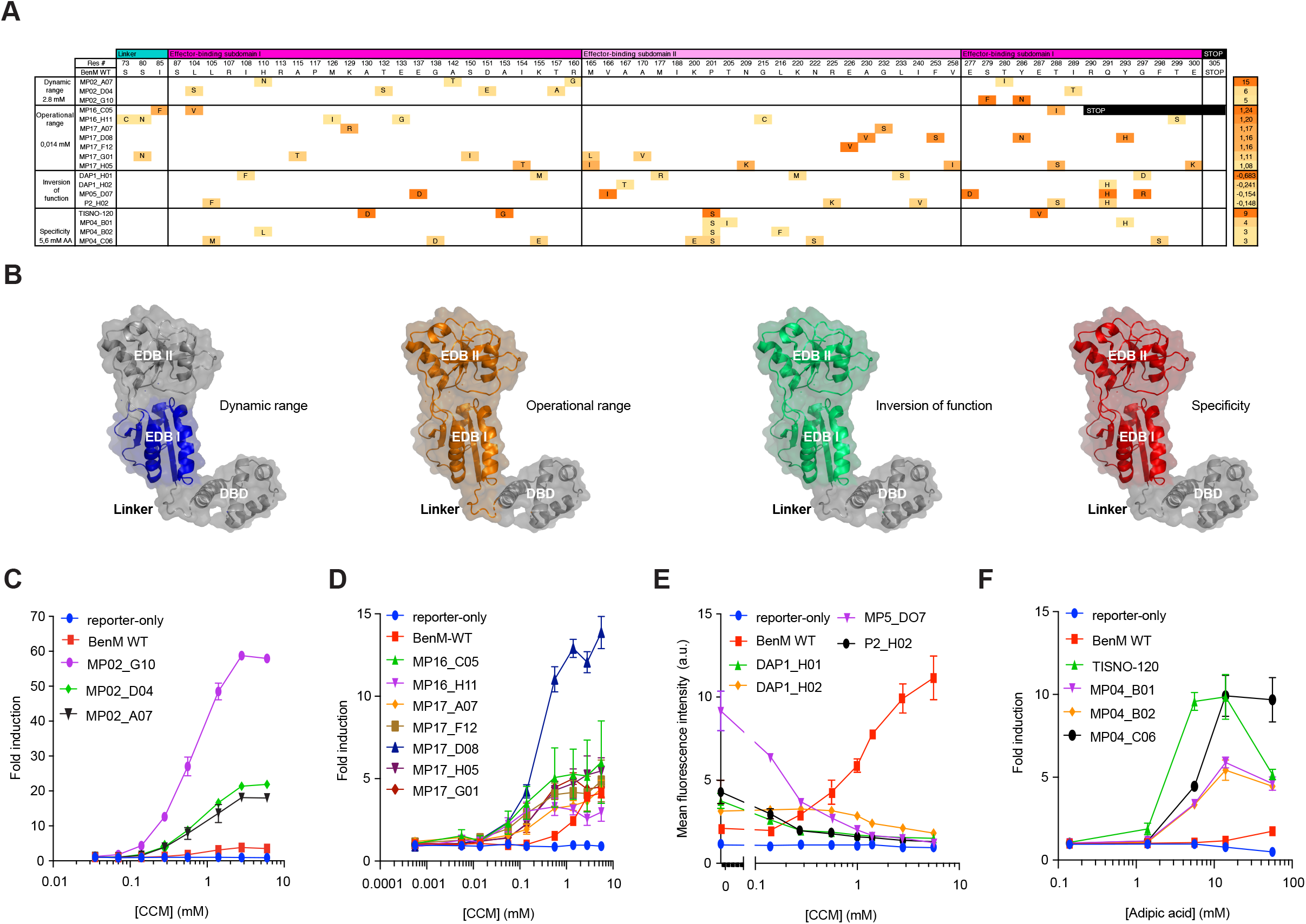
Sequence-function characterization of BenM WT and evolved archetypical variants. (**A**) Amino-acid substitution profiles for the four phenotypic categories (*Left*) of evolved BenM variants, with heat-mapping to indicate the BenM variant fold-induction for the lowest inducer concentration with a significant (*p* < 0.05) response compared to control medium. The displayed score (*Right*) for the archetypical variants of improved dynamic and operational range, as well as for change of specificity, is indicative of fold-change between inducer and control medium for the lowest inducer concentration significantly improving ON state compared to OFF state as indicated in *Methods*. For the inversion-of-function variants, the score represents the correlation coefficient between dose-response curves for WT BenM and evolved variants calculated according to equation given *Methods*. (**B**) Depictions of BenM monomeric structures (PDB 3k1n), with domains color-coded for mutational spaces for each of the four phenotypic categories. **(C-F)** Dose-response curves for BenM variants, BenM WT, and reporter-only control (No BenM). Data represents mean fold induction for dynamic range (C), operational range (D) affinity-matured (E), and fluorescence intensities for inversion-of-function variants operational range variants (F), +/- SD from three biological replicates. a.u. = arbitrary units.

For dynamic range variants we carried out a dose-response characterization of three variants (Fig. 3B). Here we found that increased CCM induction was observed at all concentrations compared to WT BenM, and that the most highly-induced variant (MP2_G10) displayed a dynamic range of 58.8 +/- 1.0, a more than 15-fold improvement over BenM WT (Fig. 3C). Notably, different from other aTF engineering efforts^43^, the increase in dynamic range was largely due to increased expression upon ligand induction, as only modest changes in OFF state expression was observed for the dynamic range variants compared to WT BenM (Fig. 2A).

For the operational range all seven titrated variants showed significant induction (*p* < 0.05) at CCM concentrations below the lower detection limit of WT BenM (i.e. 0.56 mM CCM)(Fig. 3D). Specifically, MP17_D08 significantly induced GFP expression from 0.014 mM CCM, a 40-fold improvement over WT BenM, whereas the remaining variants showed significant induction from 0.056 mM (Fig. 3D). Interestingly, whereas most variants had shifted their operational range while maintaining a similar dynamic range as WT BenM, MP17_D08 also displayed an increased dynamic range (Fig. 3D).

For the inversion-of-function variants, four variants were selected for dose-response analysis. Here, all variants showed dose-dependent deactivation for CCM (Fig. 3D), with one variant (MP05_D07) showing similar fold-change (3.9-fold reduction) as WT BenM (5-fold induction) within the operational range of WT BenM, yielding a negative correlation coefficient of *R*^2^ = −0.68 compared to the transfer function of WT BenM (Fig. 3A and Fig. 3D).

Finally, for affinity-matured BenM variants, six mutants displayed >5-fold induction by adipic acid, of which three were sequence identical to variant MP04-B01. The response curve of the four unique variants was determined for adipic acid (Fig. 3E). One variant (TiSNO120) showed significant (*p* < 0.05) induction from 1.4 mM adipic acid, whereas the three other variants were significantly induced by 5.6 mM adipic acid, whereas WT BenM was only induced from 14 mM (*p* = 0.03; Fig. 3B). With the exception of TiSNO120, the affinity-matured variants responded only modestly to CCM, corroborating the toggled selection regime (Suppl. Fig. S2, Fig. 2B).

In summary, the toggled sorting scheme of a randomly mutated aTF EBD library serves as a powerful first demonstration of the impressive evolvability of multiple aTF parameters from a single aTF platform, with aTF variants displaying up to 15-fold increased dynamic range, 40-fold change in operational range, new ligand-specificity, and inversion-of-function, compared to parental aTF.

### Structure-function relationships of evolved biosensor variants

Interestingly, several of the amino acids mutated in this study (Fig. 3A) are known to affect BenM-dependent regulation of aromatic compound catabolism in native *Acinetobacter*^44–46^, revealing the structure-function relationships governing our observed phenotypes.

Furthermore, for dynamic range mutants, mutations exclusively occur in EBD-I, while operational range variants are the only ones with mutations occurring in the linker hinge between EDB-I and the DBD (Fig. 3A and 3F). The mutational space for aTF variants with inversion of function and matured affinities for adipic acid, on the other hand, all include broad mutation windows throughout both EBD subdomains (Fig. 3F).

In order to investigate more deeply the causality and mutational space involved in aTF variant calling, it is evident from our study, that single mutant E226V in variant MP17_F12 is causal to aTF operational range. Additionally, while all specificity variants were different, they all shared the P201S mutation (Fig. 3A). Notably, residue 201 is involved in both binding of CCM and formation of the tetramer interface in BenM WT^47^, and in order to gain better understanding as to whether this residue is causal for changes in BenM ligand specificity, TiSNO120 (A130D, A153G, P201S, and E287V) was further investigated by analysing a series of single and combinatorial mutants introduced to WT BenM to evaluate their effects on biosensor read-out in response to 1.4 mM CCM or 14 mM AA (Suppl. Fig. S3). This confirmed the role of P201S in the increased response to adipic acid and decreased induction by CCM, although the re-introduction of a second mutation (E287V) was necessary to fully recapitulate the TiSNO120 phenotype.

Taken together, the sequencing and in-depth titration studies highlight regional mutational hotspots for the four different phenotypic categories of aTF variants, and further exemplifies the impact that single residue changes can have on aTF transfer function and specificity as exemplified by position E226 and P201 of BenM.

### Biochemical characterization of evolved aTF variants

Small-molecule inducible transcription factors undergo interdomain DBD-EBD allosteric transitions upon DNA and ligand binding^48^. Also, HLH-type DBD of LTTRs bind DNA as dimers or tetramers, with BenM binding to the tripartite 73-nt binding site (*benO*) in the *ben* operon of *Acinetobacter sp. ADP1* as a tetramer^49, 50^. Specifically, BenM functions as a dimer of dimers with deduced constitutive binding to dyad-symmetrical subsite 1 (ATAC-N7-GTAT), together with either subsite 2 (ATAC-N7-GTGT) or subsite 3 (ATTC-N7-GTAT), in the presence or absence of CCM, respectively (Fig. 4A-B)^50^. Interestingly, and in line with the modular architecture of aTFs, Wang *et al.* have shown that substituting the cognate operator sequence with those of homologue aTFs can impact both dynamic and operational ranges of the biosensor^51^.

**Fig. 4.**
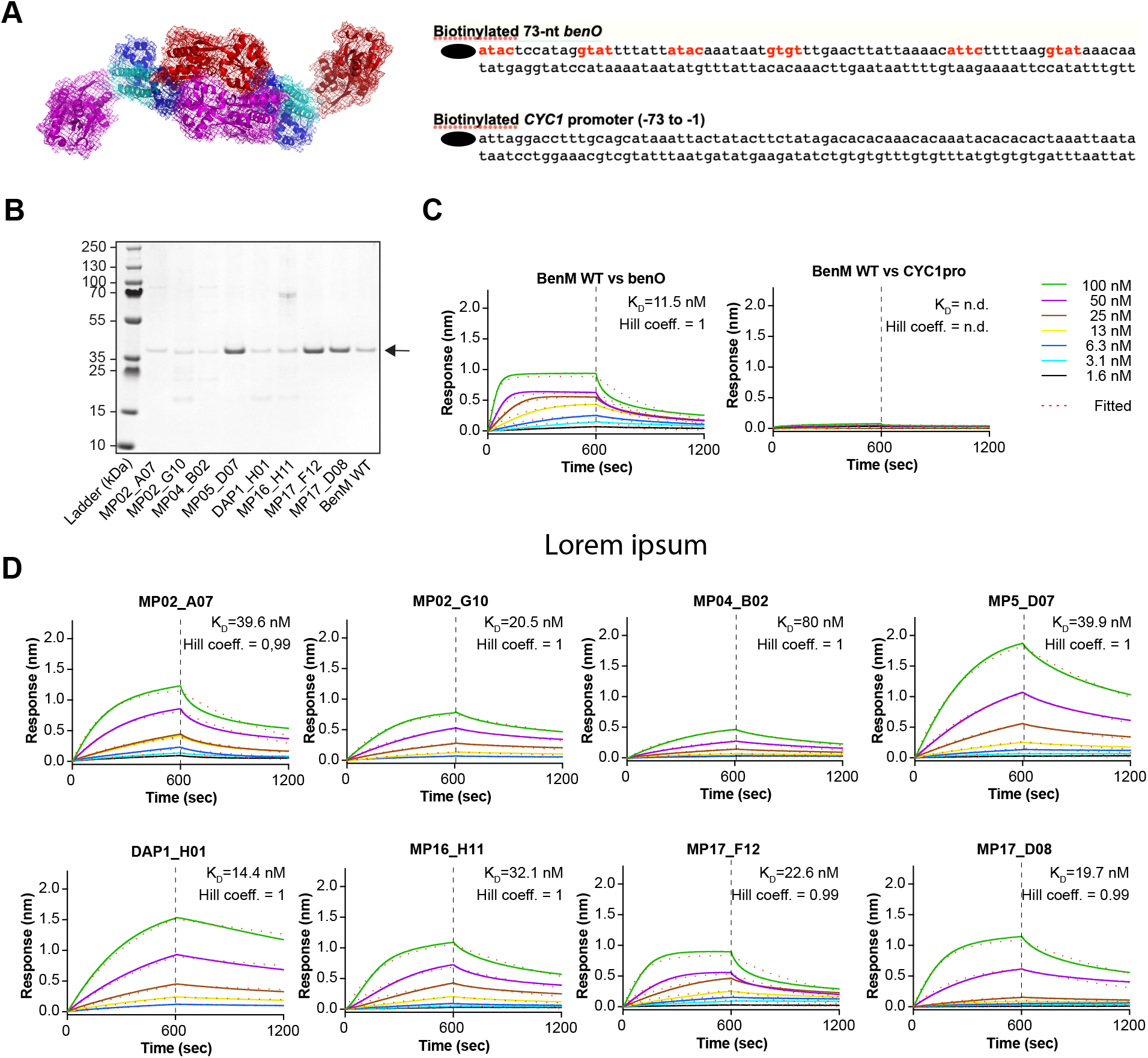
Biochemical characterization of BenM WT and evolved archetypical variants. (**A**) Tetrameric representation of full-length BenM (PDB: 3K1N) superimposed on BenM-DBD-DNA complex (PDB:4IHT)^46^. The dimeric structure of full-length BenM (PDB: 3K1N) was used to generate the probable tetrameric assembly using PDBePISA^64^ and bioassembly 1 was chosen due to the higher when compared to the alternative tetrameric bioassembly 2. The BenM-DBD-DNA complex (PDB:4IHT) was superimposed on the tetrameric assembly using PyMOL. (**B**) The sequence outline of DNA fragments 209bp_CYC1pro and 209bp_*benO*_CYC1pro, with BenM binding sites, according to Bundy *et al.*^50^, highlighted in bold red. (**C**) Expression of His_6_-tagged BenM wild type (WT) and nine archetypal variants in *E. coli* (BL21). Following batch affinity purification, the eluents from WT BenM and BenM variants were visualized on SDS/PAGE by Coomassie blue staining. Arrow indicate expected size of purified proteins. (**D**) Interaction between WT BenM WT and DNA fragments 209bp_CYC1pro and 209bp_*benO*_CYC1pro measured using biolayer interferometry (BLI) for realLtime analysis of interactions between fragments 209bp_CYC1pro and 209bp_*benO*_CYC1pro and increasing concentrations of BenM WT (1.56, 3.13, 6.25, 12.5, 25, 50 and 100LnM). Following initial loading of 100 nM biotinylated fragments 209bp_CYC1pro or 209bp_*benO*_CYC1 onto streptavidin tips, 600 sec of association and dissociation of BenM was performed (see *Methods*). The BLI signals for association at titrated BenM WT concentrations and dissociation, and the calculated *K*_d_’s are shown. (**E**) Determination of binding affinity between *benO* and BenM variants by BLI. As for (C) the BLI signals for association at 100 nM BenM variant concentrations and dissociation, and the calculated *K*_d_’s are shown. Dashed red line in (C-D) mark the fitting of the experimental association and dissociation data to a 1:1 model, with a cut-off of *R*^2^ > 0.95 of between calculated and measured binding curves for calculating binding kinetics. n.d. = not determined.

In order to determine if the transfer functions of aTF variants evolved from BenM EBD mutagenesis are coupled to changes in DNA-binding affinity, we performed biolayer interferometry (BLI). For this purpose, we heterologously expressed in *E. coli* and purified affinity-tagged WT BenM and archetypal variants from the four selection regimes (Fig. 2B-E) with homogenic purity and expected 43-44 kDa size ranges validated by SDS-PAGE^50^(Fig. 4C). Here, by titrating purified WT BenM (1.6-100 nM) onto immobilized *benO*, binding was observed, whereas no binding was observed to the negative control *cyc1* promoter element (Fig. 4D). This finding is in agreement with earlier *in vitro* studies of Bundy *et al.*^50^, and our earlier *in vivo* studies^18^, Next, performing BLI of all purified archetypical BenM variants, we also observed DNA-binding affinity for *benO* (Fig. 4D). Interestingly, evolved archetypical BenM variants had similar equilibrium dissociation constant (K_D_) for *benO* (14.3 - 80 nM) as WT BenM (11.5 nM)(Fig. 4E). In addition to K_D_’s, we calculated the Hill coefficient as described by Wang *et al.*^52^ (see *Methods*) by plotting log[Θ/(1 − Θ)] as a function of the log [BenM](Fig. 4D-E, Suppl. Fig. S6). Moreover, in line with the biochemical and structural evidence of WT BenM, all of the evolved BenM variants maintained WT BenM’s Hill coefficient of approx. 1, indicative of no cooperativity in DNA-binding (Fig. 4D-E, Suppl. Fig. S4).

Taken together, the low nM-range DNA-binding affinities observed across all archetypical BenM indicate that the evolved functionalities of the archetypal BenM variants are not due to secondary effects arising from perturbed DNA binding affinity, but rather a direct effect of EBD mutagenesis. Moreover, the elucidation of nearly identical protein structures of full-length WT BenM and two constitutively active BenM variants implies that even differences *in vivo* transcriptional activity are not well discriminated by structural studies^46^. Together these observations highlight that evolution-guided EBD designs offer a powerful method for engineering aTF functionalities, even dispensable of structural information.

### Portability of evolved biosensor variants into bacteria

Prokaryote aTFs have been ported into eukaryotes for use as biosensors for decades^53, 54^. In order to demonstrate the broader usefulness of high-throughput directed evolution of aTF-based biosensors evolved in yeast, we sought to demonstrate chassis portability. As an example biosensor we chose TiSNO120, a BenM variant affinity-matured for adipic acid biosensing - a molecule being one of the primary targets for platform chemicals in biorefineries^55^. As a recipient chassis we choose *E. coli*, a commonly used biotechnology workhorse, with adipic acid tolerance up to 50 mM (Fig. 5A)^56^.

**Fig. 5.**
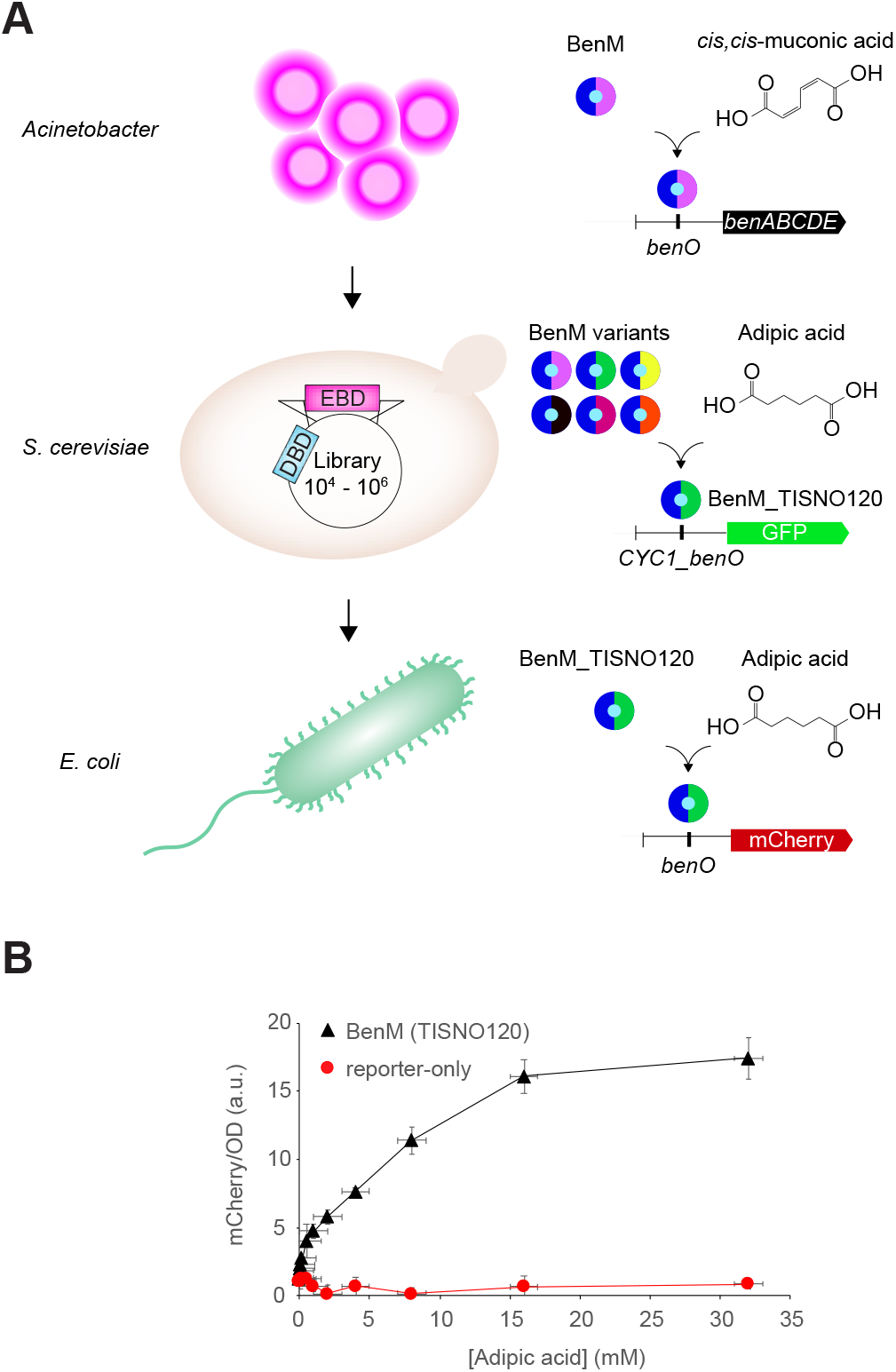
Host portability of evolved atF biosensors variants. (**A**) Schematic outline of biosensor prototyping on different chassis. (**B**) Dose-response curves for BenM variant TISNO-120 (BenM_TISNO120) affinity.-matured for adipic acid biosensing vs no BenM expression (reporter-only) in *E. coli*. Data represents mean fluorescence intensities for mCherry/OD +/- SD from three biological replicates. a.u. = arbitrary units.

Transplanting the native BenM-binding (*benO*) birectional *Acinetobacter* promoter driving the expression of TiSNO120 and mCherry into *E. coli* resulted in TiSNO120-dependent expression of mCherry within the 0-35 mM adipic acid range (Fig. 5B), exemplifying the robustness and broader usefulness of high-throughput biosensor prototyping in yeast.

## Discussion

In the present study, directed evolution coupled with toggled selection regimes was used to successfully identify novel aTF variants with user-defined transfer functions and change of specificity. To the best of our knowledge, the toggled sorting scheme of a randomly mutated aTF EBD served as a powerful first demonstration of the evolvability of multiple aTF parameters from a single aTF platform. Moreover, this study demonstrate yeast as a robust model chassis for prokaryote biosensor prototyping, and that the high-throughput workflow outlined in this study not only arms synthetic biologists with new biosensors, but also enables broader host-agnostic usability, as exemplified by the adipic acid biosensor variant being functional in *E. coli*. Finally, in this study, the starting point was BenM from *Acinetobacter sp. ADP1*, a natively CCM-inducible transcription factor, which belongs to the largest family of prokaryotic aTFs, namely LTTRs, which could as such serve as platform aTFs for further expanding the functional space of many new-to-nature biosensors.

Another important observation is the similar DNA-binding affinities broadly observed for all four classes of aTF variants (Fig. 4). For operational range variants, this indicates that beyond the strategy of using degenerate or multi-copy operator sequences to change biosensor operational ranges^51^, engineering aTFs by EBD mutagenesis is a new route to change biosensor operational range. Beyond the operational range mutants, the DNA binding affinity of inversion of function variants is particularly alluring. Given WT-like DNA binding affinity, several hypotheses on the molecular mechanisms can explain the ligand-dependent repression. Firstly, it could be speculated that the variant aTF represses rather than activates transcription by binding elsewhere in the reporter promoter, or that the mutant has increased ligand-independent DNA affinity. However, as no mutations were targeted in the DBD region of BenM, and that the BenM variants are able to bind the cognate operon for wild-type BenM with similar K_D_ (Fig. 4), both of these scenarios are considered unlikely. More likely, given the occurrence of five mutations in variant MP05_D07 (E137D, V166I, E277D, Q291H, and G297R), we hypothesize that the inversion could be related to a modification of the allosteric transition itself. In this scenario, the evolved BenM would not bind the operator with CCM bound. Instead, the variants would have a high affinity, yet be in allosteric ON state,, for the *benO* operator in the absence of CCM, analogously to the mechanism hypothesized for anti-LacI variants^30^. In addition, R225H is known to cause high OFF-state and decreased dynamic range^45^, potentially adding to the full inversion of function of variant P2_H02 containing mutation R225K in addition to four other mutations.

Although the phenotypic screen developed in this study proved useful for generating aTF variants with user-defined performance parameters, complementing biochemical characterization has also proven effective for understanding structural and mechanistic differences of mutant aTFs^57^. Since a change in phenotype can occur due to one or any combination of (i) altered specificity and/or affinity for an inducer, (ii) expression levels and protein stability, (iii) DNA binding affinity, and (iv) presence of protein subunits in active or inactive conformations, the more detailed mechanistic effects should be investigated for purified aTF variants from different phenotypic categories^57^. For instance, for any oligomeric DNA-binding aTF, it will be relevant to analyze the oligomerization state and molecular dynamics, as well as ligand affinity to determine how each of these functions could affect aTF variant performance, and thus improve our understanding of how individual or combinations of mutations affect aTF structure and function, and thereby enable more rational biosensor engineering.

Ultimately, combining the functional and high-throughput evolution-guided method presented in this study, together with computational design of ligand binding and structural dynamics, should enable a multi-faceted approach, necessary to gain better understanding of biosensor allostery, and how this can be forward-engineered for future designer biosensors.

## Methods

### Medium and chemicals

Stable *S. cerevisiae* strains were routinely cultivated at 30 °C on YPD (1% (w/v) yeast extract, 2% (w/v) peptone 2% (w/v) dextrose, solidified with 2% (w/v) agar) medium, whereas plasmid-containing strains or libraries were cultivated in synthetic complete medium lacking leucine (SC-Leu; 6.7 g/L yeast nitrogen base without amino acids, 1.62 g/L yeast synthetic drop-out medium supplement without leucine (Y1376, Sigma-Aldrich), 2% (w/v) dextrose, pH 5.6, 2% (w/v) agar in case of plates). Mineral medium supplemented with tryptophan (7.5 g/L (NH_4_)_2_SO_4_, 14.4 g/L KH_2_PO_4_, 0.5 g/L MgSO_4_•7H_2_O, 2 g/L dextrose, trace metals, vitamins, 0.02 g/L tryptophan, pH 4.5) was freshly prepared as described previously^58^, and medium was handled as outlined by Ambri *et al.*^59^. Diacids, *cis, cis*-muconic acid (Sigma, CAS Number 1119-72-8, Product Number: 15992) or adipic acid (TCI, CAS Number:124-04-9 Product Number: A0161), were dissolved freshly to the medium on the day of handling, after which the pH was adjusted to 4.5 and the solution filter-sterilized as previously described^59^.

EasyClone plasmids used in this paper are outlined in Jensen *et al*.^58^. *Escherichia coli strain* DH5 was used as a host for cloning and plasmid propagation, and cultivated at 37 °C in Luria-Bertani (LB) medium supplemented with 100 ug/mL ampicillin. PCR was carried out using Phusion® or Phusion U High-Fidelity DNA Polymerase (New England Biolabs) according manufacturer’s instructions. Site-directed mutagenesis of BenM in pMeLS0076 was carried out using the QuikChange Site-Directed Mutagenesis Kit (Agilent Technologies) according to manufacturer’s instructions.

Yeast transformation was performed according to Gietz & Schiestl^60^, followed by selection on synthetic drop-out medium.

All oligonucleotides, plasmids, strains, and synthetic DNA used in this study are listed in Suppl. Table S2.

### Library generation

In order to generate a library of mutated fragments of the BenM-EBD the procedure described by Skjoedt *et al*.^18^ was followed. In short, epPCR was carried out using the Agilent GeneMorph mutagenesis II kit following manufacturer’s instructions for high mutational load. Five consecutive rounds of epPCR were carried out using primers MelS69-F and MelS93-R, using 50 ng pMelS0076 as a starting template for the first round of PCR. After each round, the 735-bp band was gel-purified, and 50 ng was used as input for the next round. To make a library of yeast strains, mutated fragments from rounds 2-5 were used as input for four regular PCRs (Phusion) using tailed primers MelS071-F and MelS094-R and column-purified. Pooled fragments from rounds 2 and 3 (sublibrary 1), as well as rounds 4 and 5 (sublibrary 2), were co-transformed with gapped vector pMelS0076 into strain MelS009 (CEN.PK with chromosomal integration of yeGFP reporter gene) in a molar ratio of 80:1, in order to allow for re-constitution of the plasmid. For each sublibrary two transformations were carried out (4×10^7^ cells as input each), the total biomass was pooled afterwards, added to 48 mL SC-leu, grown overnight to saturation and frozen in aliquots (sublibrary 1, TISNO-122; sublibrary 2, TISNO-123). From plating for single colonies right after transformation it was found that sublibrary 1 contained 224,000 variants, and sublibrary 2 contained 198,000 variants. Colony PCR and sequencing of 10 random colonies per sublibrary narrowed down the library size for sublibrary 1 to 45,000 effective variants, and sublibrary 2 to 40,000 effective variants. In this article sublibrary 1 and 2 are always combined for sorting purposes (effectively containing 85 000 variants) and referred to as ‘the BenM library’.

### FACS-based selection

For all sorting steps *S. cerevisiae* cells containing the BenM library or subpopulations thereof previously frozen in 25% (v/v) glycerol at −80 °C were thawed and added to 5 mL SC-Leu to become OD_600_ = 0.2 (approx. 2 × 10^6^ cells/mL). Cultures were grown overnight in 12-mL preculture tubes in a 30 °C incubator shaking at 300 rpm with 5 cm orbit diameter. The next day, cultures were diluted into minimal medium +/- inducer to OD_600_=0.2 and incubated under the same conditions. In parallel, control strains were processed similarly. After 22 h each culture was diluted in 2 mL sterile 1x PBS to an OD_600_ of approx. 1.0 right before being analyzed on a Becton Dickinson Aria fluorescence-assisted cell sorting (FACS) instrument with a blue laser (488 nm) to detect yeGFP fluorescence. For control strains 10,000 single-cell events were measured and for (sub)libraries 250,000 events were measured to assess mean fluorescence intensity and diversity of each population in medium with or without inducer. Depending on the phenotype under study, specific gates were drawn to select for single-cell events obeying the set criterion (Fig. S2). In any case, cells were sorted in FITC-A vs FSC-A pot in order to select based on fluorescence intensity without bias for a certain cell size as carried out previously^61, 62^. Sorted cells were collected in 2-5 mL SC-Leu (initial cell density < 5,000 cells/ mL) and allowed to recover overnight (30 °C, 300 rpm) until a sufficient OD_600_ was reached after which cells were frozen down in 25% (v/v) glycerol at −80 °C in aliquots containing 2.5 OD_600_ units each.

### Flow cytometry

Flow cytometry screening was carried out as described previously^18^. In short, clonal variants were inoculated into 150 μL SC-Leu in 96-well plates and grown overnight. The next day, strains were subcultured 1:100 into 500 μL medium +/- inducer in a deep-well plate and cultured for 22-23 h before analyzed by flow cytometry. In parallel, control strains ST.1 (WT CEN.PK), MeLS0138 (reporter-only) and MeLS0275 (BenM WT – reporter) were processed in the same fashion in biological triplicates in each experiment. The mean fluorescence intensity (MFI) of each strain was determined based on 10 000 gated single-cell events. The normalized OFF state of each variant was determined by dividing its MFI in control medium by the average MFI of MeLS0275 in control medium in the same experiment.

### Protein expression and purification

BenM and its variants were expressed in *E. coli* BL21 (DE3). An overnight culture grown in 2xYT was used to inoculate 0.5 L MagicMedia (Invitrogen), both supplemented with 50 µg/mL kanamycin, grown at 37 °C and shaking at 200 rpm. When the OD_600_ reached 0.6, cells were transferred to 18 °C and incubated for 36 hours. After harvest, cells were lysed by three passes through an emulsiflex C5 (Avestin, Mannheim, Germany) and cleared cell lysate was loaded onto a HisTrap FF column (GE Healthcare). After washing with 10 column volumes of buffer A (30 mM Tris-HCl, 500 mM NaCl, 30% Glycerol, 5 mM Imidazole, 1 mM DTT pH 7.9), proteins were eluted with 4 column volumes each of 5%, 10%, 15% and 100% buffer B (30 mM Tris-HCl, 500 mM NaCl, 30% Glycerol, 500 mM Imidazole, 1 mM DTT pH 7.9). When purity was unsatisfactory, protein were polished on a HiLoad 16/600 Superdex 200pg column equilibrated in 50 mM Tris-HCl, 150 mM NaCl pH 8.0. Proteins were frozen in N_2_(l) and stored at −80 °C.

### Biolayer interferometry

Synthetic 5’-end biotinylated (forward) and non-modified (reverse) single-stranded DNA oligos were ordered from IDT (Integrated DNA Technologies, Coralville, IA), and resuspended to final concentrations of 200 nM in IDT Nuclease-free duplex buffer according to manufacturer’s instructions. Complementary single-stranded oligos for preparing 209bp_CYC1p (TISNO-110 and TISNO-113) and 209bp_*benO*_CYC1p (TISNO-112 and TISNO-114) were combined in 1:1 ratios using 20 uL of each oligo, and incubated at 94 °C for 2 min. Next the mixtures were left at RT for 3 hrs before storing them as 2.5 uL aliquots of 100 μM DNA (0.25 mol/aliquot). Using 0.2 nmol of DNA for each experiment, all DNA binding experiments were performed using an FortéBio Octet RED96 (Pall, Menlo Park, CA, USA) mounted with black 96-well microplates at 30 °C with 200 μL volume.

Streptavidin sensors (Pall, Menlo Park, CA, USA) were pre-equilibrated in PBS buffer for 600s and loaded with biotinylated 209bp_CYC1pro or 209bp_*benO*_CYC1pro (100 nM, 600s). After reaching baseline for 300s, association and dissociation of the indicated concentrations of BenM WT and variants were measured for 600s each. All assay steps were performed in kinetics buffer (PBS containing 0.02% Tween-20 v/v and 0.1% BSA w/v, pH 7.4). Binding kinetics were calculated using the FortéBio Data Analysis v7.1 software by fitting the association and dissociation data to a 1:1 model. A cut-off of *R*^2^ > 0.95 was used in order to ensure binding kinetics were calculated only when there was a good fit between calculated and measured binding curves.

The measured signal at equilibrium, noted Req, and the calculated Rmax for each protein concentration were used to calculate fractional saturation and plotted against the respective protein concentration to determine the Hill coefficient according to Engohang-Ndong *et al.*^63^

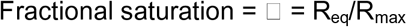

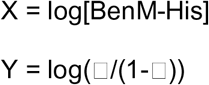

### Analysis of BenM variants by sequencing

Fold-change (FC) fluorescence induction was calculated for each data point comparing the fluorescence level in the control medium and the fluorescence level in the inducer media. The displayed score for the phenotypes of improved dynamic and operational range, as well as for change of specificity, were computed normalising the mutants’ FC values by the FC value of the wild type protein at the same conditions. The Pearson correlation coefficient (PCC) was calculated using FC values of the mutated phenotype and FC values of the wild type phenotype at increasing concentrations. The dynamic ranges were calculated for the wild type (R_wt_) and for the mutant (R_v_) phenotypes, as the difference between the corresponding highest and lower values of FC:

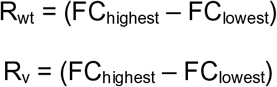

Additionally, the ratio between R_v_ and R_wt_ was used as coefficient to indicate the similarity of dynamic range between the mutant and the wild type. The inversion of function phenotype score (X) was calculated multiplying the PCC and the ratio between R_v_ and R_wt_:

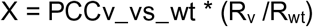

Where the highest inversion of function corresponds to high anti-correlation (PCC ∼ −1) and similar dynamic range (R_v_ /R_wt_ ∼1).

### Biosensing in E. coli

The adipic acid biosensing plasmid (JBx_101898) was assembled of 4 DNA parts by NEBuilder® HiFi DNA Assembly Master Mix (New England Biolabs, MA): First, the 3.3 kb backbone was amplified from pBbS5c-RFP (JBp_000017) using primers j5_00193_(pBbS5c_backbone)_forward and j5_00194_(pBbS5c_backbone)_reverse. Next, the 0.9 kb *benM_Mu1* was amplified from pTS-56 using primers j5_00195_(BenM_Mu_rc)_forward and j5_00196_(BenM_Mu_rc)_reverse, while BP-01, the 0.3 kb DNA binding site of BenM (*benO*) with overlap regions, was synthesized as gBlocks® Gene Fragment (Integrated DNA Technologies, Inc., IA). Lastly, the 0.7 kb *mCherry* was amplified from pSC-gapdhp(EL)-mCherry (JBx_065530) using primers j5_00198_(mCherry)_forward and j5_00199_(mCherry)_reverse. The negative control plasmid (JBx_101899) was constructed via self-circularization: primers BenM_KO_R and BenM_KO_F were used to amplify the 4.3 kb fragment from JBx_101898, the PCR product was self-circularized by T4 ligase (Thermo Fisher Scientific Inc., MA).

The *E. coli* DH10B transformed with JBx_101898 (JBx_101900) was used for adipic acid biosensing, while the *E. coli* DH10B transformed with JBx_101899 (JBx_101901) was used as negative control. The *E. coli* cell was grown overnight in Luria−Bertani (LB) medium containing 25 μg/mL chloramphenicol at 37 °C. The overnight culture was inoculated 1:100 at 50 mL fresh LB medium containing chloramphenicol and grown at 37 °C. Once the OD600 reached 0.6, the culture was cooled down to room temperature and aliquoted into 96-well deep well plate (2 mL volume, V-bottom) at 0.5 mL per well. The adipic acid aqueous stock was prepared at 1.7 M with pH = 7.0 (adjusted by NaOH) and filtered with 0.22 μm filter. The working solutions with varying concentrations were diluted with sterile water from the stock. 10 μL of working solution was added to each well to make the adipic acid final concentrations (mM) at 0, 0.02, 0.05, 0.1, 0.2, 0.5, 1, 2, 4, 8, 16, 32. The deep well plate was sealed by AeraSeal film (Omega Bio-tek, Inc., GA) and wrapped by aluminum foil. The plate was shaken at 250 rpm in 30 °C for 16 h. After that, 100 μL culture from each well was transferred into 96-well Costar® assay plate (black with clear flat bottom, Corning Inc., NY) to measure the optical density at the wavelength of 600 nm absorbance and mCherry fluorescence (λex = 575 nm, λem = 620 nm, λcutoff = 610 nm) by SpectraMax M2 plate reader (Molecular Devices, LLC., CA). The assays were performed in triplicate wells for each concentration.

## Acknowledgements

This work was supported by the Novo Nordisk Foundation.

## Author contributions

T.S., E.K.C., J.D.K., and M.K.J. conceived this project. T.S., E.K.C., B.P., J.F.B., S. P.B., and S.K. designed all of the experiments. F.A., E.K.C., and D.H.W performed all structure-function analysis. T.S., E.K.C., B.P., J.F.B., S. P.B., S.K. and M.K.J analyzed the data. T.S. and M.K.J. wrote the paper.

## Supplementary material

**Supplementary Fig. S1. Multiple sequence alignment of the protein sequences of 12 LTTRs.** Alignment, including WT BenM and other yeast-implemented LTTRs FdeR, ArgP, MdcR and PcaQ, was generated using T-Coffee and visualized using Boxshade^65^. Fraction cut-off for sequence shading was set at 0.7.

**Supplementary Fig. S2. CCM-responsiveness of adipic acid affinity-matured BenM variants.** Measurements are means +/- SEM from three independent biological replicates. a.u. = arbitrary units.

**Supplementary Fig. S3. Effects of single and double mutations in biosensor performance.** Individual and combinations of mutations found in BenM variant TiSNO120 were introduced to WT BenM, and transformed into a reporter strain expressing GFP under the control of the truncated synthetic 209_CYC1_*benO* promoter in yeast. CCM = *cis,cis*-muconic acid. AA = Adipic acid. a.u. = arbitrary units.

**Supplementary Fig. S4. Hill plot for WT BenM and BenM variants binding to DNA fragment *benO*.** The equation is indicated for each protein and the slope of 0.99-1.00 corresponds to the Hill coefficient. Req was measured and Rmax was calculated independently for each protein concentration. The calculations were performed as previously described^63^.

**Supplementary Table S1.** Overview of FACS regimes for evolution-guided aTF transfer function engineering and affinity-maturation.

**Supplementary Table S2.** Overview of strain, plasmids, primers, and gBlocks used in this study.

