## Supplementary information for "Evolution-guided engineering of small-molecule biosensors"

### Supplementary material

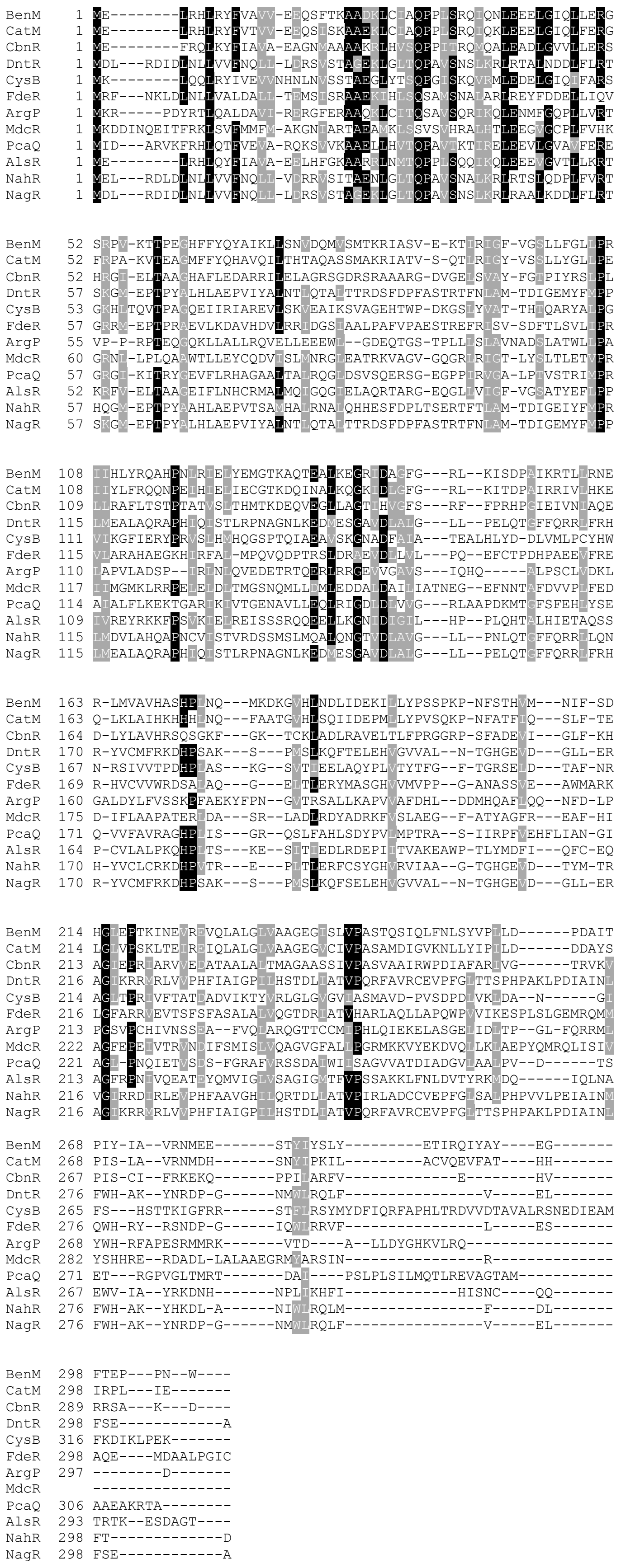

**Supplementary Figure S1. Multiple sequence alignment of the protein sequences of 12 LTTRs.** Alignment, including WT BenM and other yeast-implemented LTTRs FdeR, ArgP, MdcR and PcaQ, was generated using T-Coffee and visualized using Boxshade[65]. Fraction cut-off for sequence shading was set at 0.7.

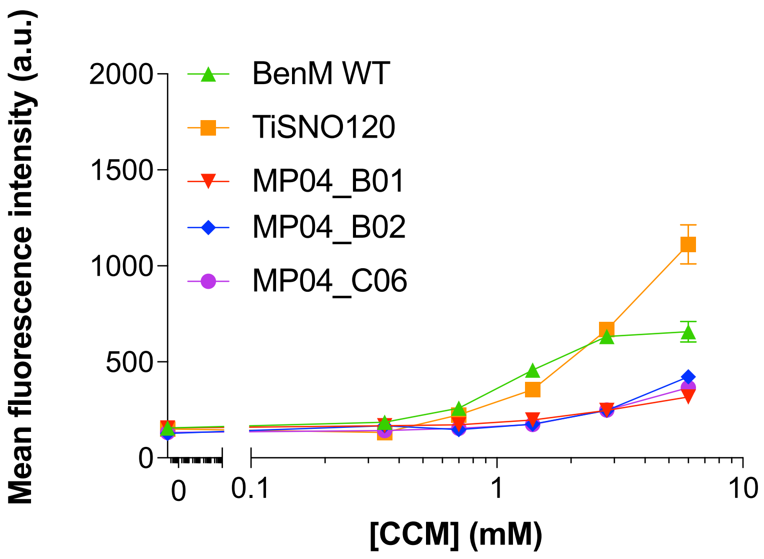


**Supplementary Fig. S2. CCM-responsiveness of adipic acid affinity-matured BenM variants.** Measurements are means +/- SEM from three independent biological replicates. a.u. = arbitrary units.

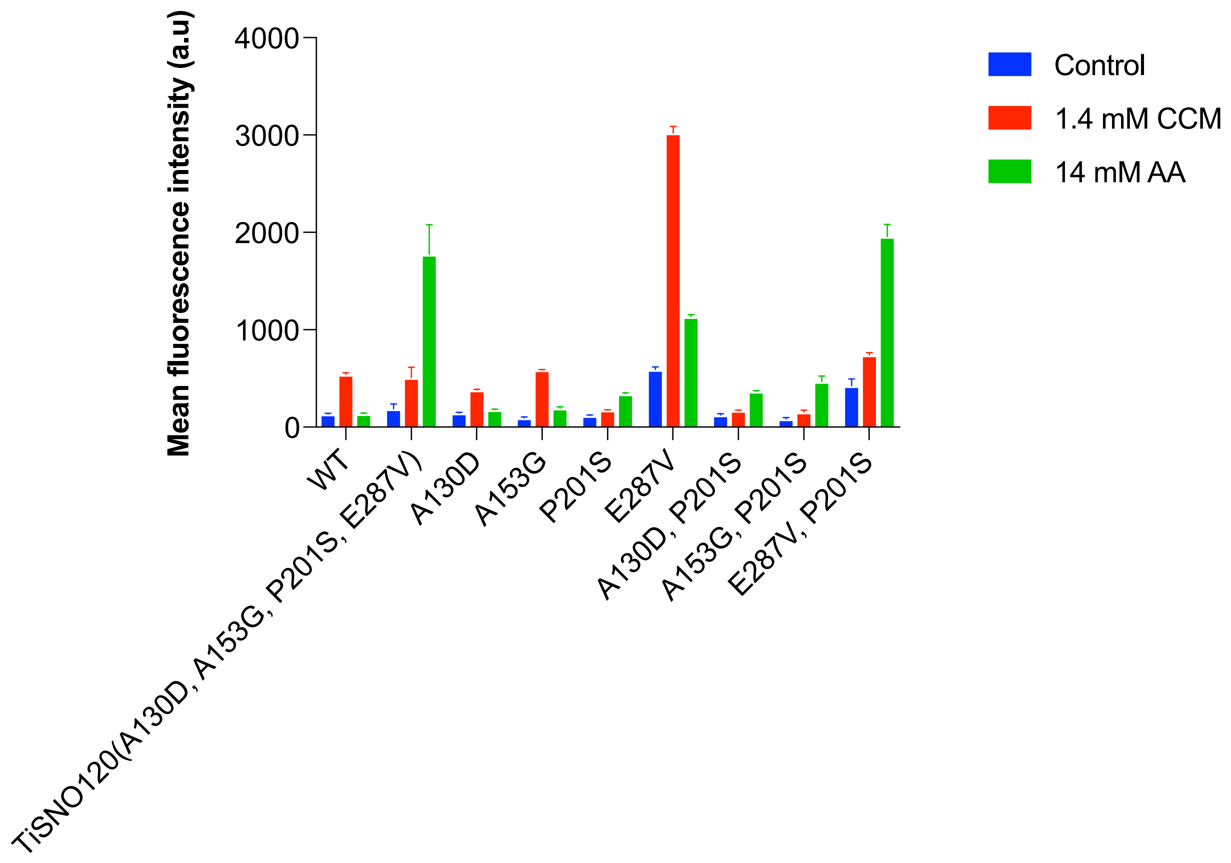


**Supplementary Fig. S3. Effects of single and double mutations on biosensor performance.** Individual and combinations of mutations found in BenM variant TiSNO120 were introduced to WT BenM. While E287 increased OFF state and induction by both CCM and AA, single mutant P201S decreased CCM induction and increased AA induction. Double mutant P201S, E287V showed a phenotype similar to quadruple mutant TiSNO120, although with increased background, and induced expression levels. Error bars show standard deviation based from three replicate measurements. CCM = *cis,cis*-muconic acid. AA = adipic acid. a.u. = arbitrary units.

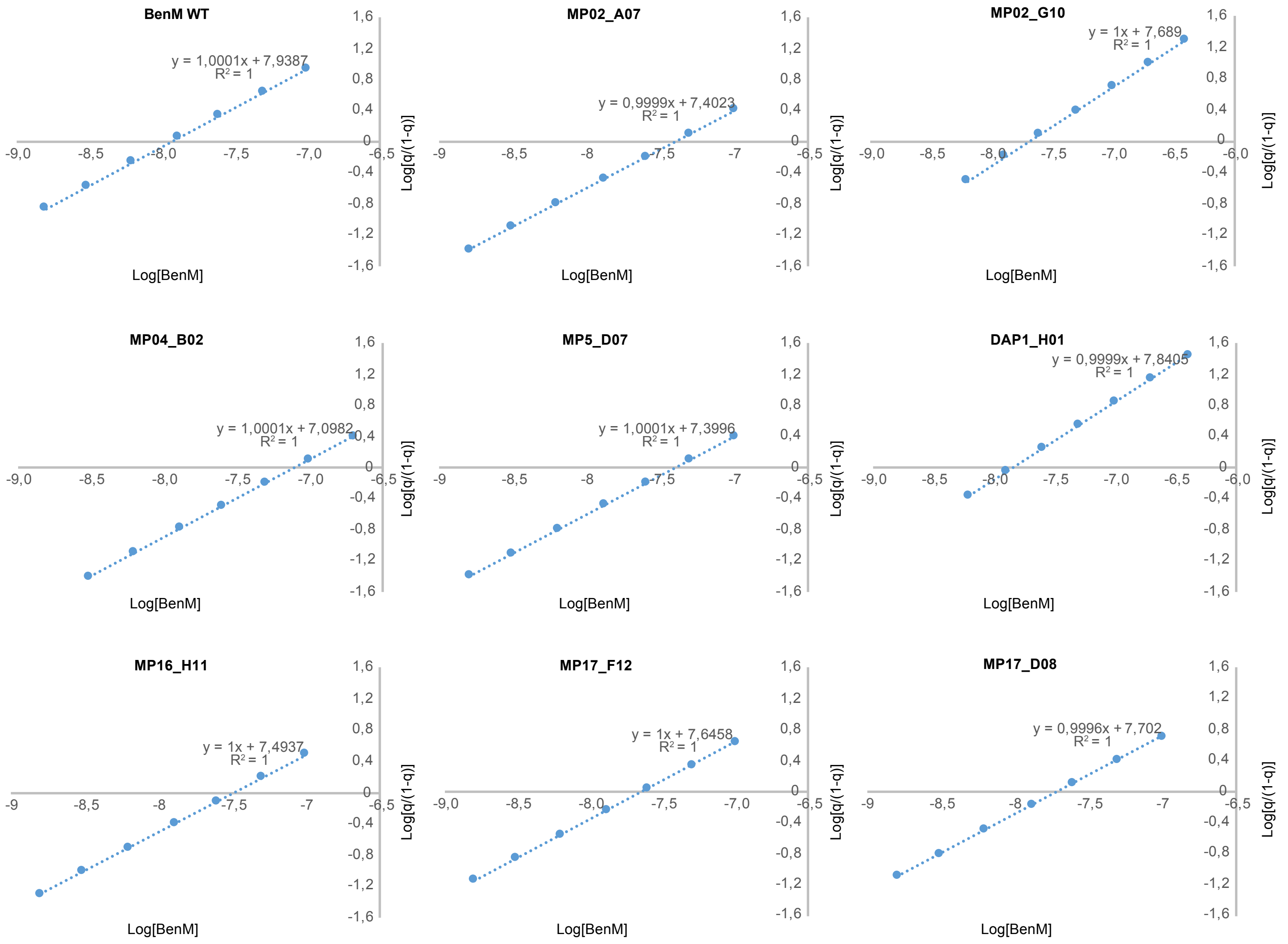


**Supplementary Fig. S4. Hill plot for WT BenM and BenM variant binding to DNA fragment benO.** The equation is indicated for each protein and the slope of 0.99-1.00 corresponds to the Hill coefficient. $R_{eq}$ was measured and $R_{max}$ was calculated independently for each protein concentration. The calculations were performed as previously described[63].

**Supplementary Table S1.** Overview of FACS regimes for evolution-guided aTF transfer function engineering and change of specificity.

| | | Selection I | | | Selection II |
|---|---|---|---|---|---|
| **Dynamic range** | Exp. #116 | | | Exp. #116 | |
| | Sorted population | TISNO-122+123 | | Sorted population | TISNO-134 |
| | Sorted media, mode | 1.4 mM CCM, ON | | Sorted media, mode | Delft, OFF |
| | Gate based on | TISNO-122+123, Delft | | Gate based on | MelS0138, Delft |
| | %sorted | 0.17 | | %sorted | 43.4 |
| | # variants sorted (non-specific variants) | 144 (14) | | # variants sorted (non-specific variants) | 62 (36) |
| **Change of specificity** | Exp. #116 | | | Exp. #116 | |
| | Sorted population | TISNO-122+123 | | Sorted population | TISNO-135 |
| | Sorted media | 14 mM AA, ON | | Sorted media, mode | Delft, OFF |
| | Gate based on | TISNO-122+123, Delft | | Gate based on | MelS0138, Delft |
| | %sorted | 0.032 | | %sorted | 40.5 |
| | # variants sorted (non-specific variants) | 27 (14) | | # variants sorted (non-specific variants) | 11 (3) |
| **Operational range** | Exp. #164 | | | Exp. #164 | |
| | Sorted population | TISNO-122+123 | | Sorted population | TISNO-122+123 |
| | Sorted media | 35 uM, ON | | Sorted media | 70 uM, ON |
| | Gate based on | top-1% | | Gate based on | top-1% |
| | %sorted | 1.13 | | %sorted | 1.12 |
| | # variants sorted (non-specific variants) | 961 (816) | | # variants sorted (non-specific variants) | 952 (765) |
| **Inversion of function** | Exp. #116 | | | Exp. #178 | |
| | Sorted population | TISNO-122+123 | | Sorted population | TISNO-138 |
| | Sorted media | Delft, ON | | Sorted media | 5.6 mM CCM, OFF |
| | Gate based on | MelS0138, Delft | | Gate based on | MelS0138, Delft |
| | %sorted | 5.49 | | %sorted | 34.9 |
| | # variants sorted (non-specific variants) | 4667 (8262) | | # variants sorted (non-specific variants) | 1629(3323) |

**Supplementary Table S2.** Overview of strain, plasmids, primers, and synthetic DNA used in this study.

| Strain name | Description | Plasmid/ modification (parent strain) | Genotype | Source |
|---|---|---|---|---|
| CEN.PK113-7D | Prototrophic WT *S. cerevisiae* strain | - | mat a URA3 HIS3 LEU2 TRP1 | Peter Kötter |
| MeLS0009 | Donor strain library | - | mat a his3 leu2 trp1 209bp_CYC1p_BenO_T1::yEGFP-SpHIS5 | Skjoedt et al, 2016[1] |
| MeLS0138 | Reporter-only control strain | - | mat a his3 leu2 trp1 KI.LEU 209bp_CYC1p_BenO_T1::yEGFP-SpHIS5 | Skjoedt et al, 2016[1] |
| MeLS0275 | Plasmid-based BenM WT + reporter strain | - | mat a his3 leu2 trp1 + REV1p::BenM-KI.LEU2 209bp_CYC1p_BenO_T1::yEGFP-SpHIS5 | Skjoedt et al, 2016[1] |
| TISNO-120 | Variant TISNO-120 re-transformed (specificity) | pTS-56 (MeLS0009) | mat a his3 leu2 trp1 + REV1p::BenM$^{TISNO-120}$-KI.LEU2 209bp_CYC1p_BenO_T1::yEGFP-SpHIS5 | This study |
| TISNO-122 | Library derived from second and third round of epPCR on BenM-EBD | pMeLS0076* (MeLS0009) | mat a his3 leu2 trp1 + REV1p::BenM*-KI.LEU2 209bp_CYC1p_BenO_T1::yEGFP-SpHIS5 | This study |
| TISNO-123 | Library derived from fourth and fifth round of epPCR on BenM-EBD | pMeLS0076* (MeLS0009) | mat a his3 leu2 trp1 + REV1p::BenM*-KI.LEU2 209bp_CYC1p_BenO_T1::yEGFP-SpHIS5 | This study |
| TISNO-155 | Variant BenM(P201S) | pTS-66 (MeLS0009) | mat a his3 leu2 trp1 + REV1p::BenM(P201S)-KI.LEU2 209bp_CYC1p_BenO_T1::yEGFP-SpHIS5 | This study |
| TISNO-158 | Variant MP02_A7 isolated by FACS (dynamic range) | FACS isolate (TISNO-122 + 123) | mat a his3 leu2 trp1 + REV1p::BenM$^{MP02_A07}$-KI.LEU2 209bp_CYC1p_BenO_T1::yEGFP-SpHIS5 | This study |
| TISNO-159 | Variant MP02_D04 isolated by FACS (dynamic range) | FACS isolate (TISNO-122 + 123) | mat a his3 leu2 trp1 + REV1p::BenM$^{MP02_D04}$-KI.LEU2 209bp_CYC1p_BenO_T1::yEGFP-SpHIS5 | This study |
| TISNO-160 | Variant MP04_C06 isolated by FACS (specificity) | FACS isolate (TISNO-122 + 123) | mat a his3 leu2 trp1 + REV1p::BenM$^{MP04_C06}$-KI.LEU2 209bp_CYC1p_BenO_T1::yEGFP-SpHIS5 | This study |
| TISNO-161 | Variant MP05_D07 isolated by FACS (inversion of function) | FACS isolate (TISNO-122 + 123) | mat a his3 leu2 trp1 + REV1p::BenM$^{MP05_D07}$-KI.LEU2 209bp_CYC1p_BenO_T1::yEGFP-SpHIS5 | This study |

| | | | | |
|---|---|---|---|---|
| TISNO-185 | Variant BenM(A130D) | pTS-92 (MeLS0009) | mat a his3 leu2 trp1 + REV1p::BenM(A130D)-KI.LEU2 209bp_CYC1p_BenO_T1::yEGFP-SpHIS5 | This study |
| TISNO-186 | Variant BenM(A153G) | pTS-93 (MeLS0009) | mat a his3 leu2 trp1 + REV1p::BenM(A153G)-KI.LEU2 209bp_CYC1p_BenO_T1::yEGFP-SpHIS5 | This study |
| TISNO-187 | Variant BenM(E287V) | pTS-94 (MeLS0009) | mat a his3 leu2 trp1 + REV1p::BenM(E287V)-KI.LEU2 209bp_CYC1p_BenO_T1::yEGFP-SpHIS5 | This study |
| TISNO-188 | Variant BenM(P201S, A130D) | pTS-95 (MeLS0009) | mat a his3 leu2 trp1 + REV1p::BenM(A130D, P201S)-KI.LEU2 209bp_CYC1p_BenO_T1::yEGFP-SpHIS5 | This study |
| TISNO-189 | Variant BenM(P201S, A153G) | pTS-96 (MeLS0009) | mat a his3 leu2 trp1 + REV1p::BenM(A153G, P201S)-KI.LEU2 209bp_CYC1p_BenO_T1::yEGFP-SpHIS5 | This study |
| TISNO-190 | Variant BenM(P201S, E287V) | pTS-97 (MeLS0009) | mat a his3 leu2 trp1 + REV1p::BenM(P201S, E287V)-KI.LEU2 209bp_CYC1p_BenO_T1::yEGFP-SpHIS5 | This study |
| TISNO-193 | Variant MP02_A7 re-transformed (dynamic range) | pTS-100 (MeLS0009) | mat a his3 leu2 trp1 + REV1p::BenM(H110N, A142T, R160G, T280I)-KI.LEU2 209bp_CYC1p_BenO_T1::yEGFP-SpHIS5 | This study |
| TISNO-194 | Variant MP04_C06 re-transformed (specificity) | pTS-101 (MeLS0009) | mat a his3 leu2 trp1 + REV1p::BenM(L105M, G138D, K155E, K200E, P201S, N222S, F298S)-KI.LEU2 209bp_CYC1p_BenO_T1::yEGFP-SpHIS5 | This study |
| TISNO-195 | Variant MP05_D07 re-transformed (inversion of function) | pTS-102 (MeLS0009) | mat a his3 leu2 trp1 + REV1p::BenM(E137D, V166I, E277D, Q291H, G297R)-KI.LEU2 209bp_CYC1p_BenO_T1::yEGFP-SpHIS5 | This study |
| TISNO-196 | Variant MP02_G10 isolated by FACS (dynamic range) | FACS isolate (TISNO-122 + 123) | mat a his3 leu2 trp1 + REV1p::BenM$^{MP02_G10}$-KI.LEU2 209bp_CYC1p_BenO_T1::yEGFP-SpHIS5 | This study |
| TISNO-213 | Variant P2_H02 isolated by FACS (inversion of function) | FACS isolate (TISNO-122 + 123) | mat a his3 leu2 trp1 + REV1p::BenM$^{P2_H02}$-KI.LEU2 209bp_CYC1p_BenO_T1::yEGFP-SpHIS5 | This study |
| TISNO-214 | Variant DAP1_H01 isolated by FACS (inversion of function) | FACS isolate (TISNO-122 + 123) | mat a his3 leu2 trp1 + REV1p::BenM$^{DAP1_H01}$-KI.LEU2 209bp_CYC1p_BenO_T1::yEGFP-SpHIS5 | This study |

| | | | | |
|---|---|---|---|---|
| TISNO-215 | Variant DAP1_H02 isolated by FACS (inversion of function) | FACS isolate (TISNO-122 + 123) | mat a his3 leu2 trp1 + REV1p::BenM$^{DAP1_H02}$-KI .LEU2 209bp_CYC1p_BenO_ T1::yEGFP-SpHIS5 | This study |
| TISNO-224 | Variant MP16_C05 isolated by FACS (operational range) | FACS isolate (TISNO-122 + 123) | mat a his3 leu2 trp1 + REV1p::BenM$^{MP16_C05}$-KI .LEU2 209bp_CYC1p_BenO_ T1::yEGFP-SpHIS5 | This study |
| TISNO-225 | Variant MP16_H11 isolated by FACS (operational range) | FACS isolate (TISNO-122 + 123) | mat a his3 leu2 trp1 + REV1p::BenM$^{MP16_H11}$-KI .LEU2 209bp_CYC1p_BenO_ T1::yEGFP-SpHIS5 | This study |
| TISNO-226 | Variant MP17_A07 isolated by FACS (operational range) | FACS isolate (TISNO-122 + 123) | mat a his3 leu2 trp1 + REV1p::BenM$^{MP17_A07}$-KI .LEU2 209bp_CYC1p_BenO_ T1::yEGFP-SpHIS5 | This study |
| TISNO-228 | Variant MP17_F12 isolated by FACS (operational range) | FACS isolate (TISNO-122 + 123) | mat a his3 leu2 trp1 + REV1p::BenM$^{MP17_F12}$-KI .LEU2 209bp_CYC1p_BenO_ T1::yEGFP-SpHIS5 | This study |
| TISNO-229 | Variant MP17_D08 isolated by FACS (operational range) | FACS isolate (TISNO-122 + 123) | mat a his3 leu2 trp1 + REV1p::BenM$^{MP17_D08}$-KI .LEU2 209bp_CYC1p_BenO_ T1::yEGFP-SpHIS5 | This study |
| TISNO-230 | Variant MP17_G01 isolated by FACS (operational range) | FACS isolate (TISNO-122 + 123) | mat a his3 leu2 trp1 + REV1p::BenM$^{MP17_G01}$-KI .LEU2 209bp_CYC1p_BenO_ T1::yEGFP-SpHIS5 | This study |
| TISNO-231 | Variant MP17_H05 isolated by FACS (operational range) | FACS isolate (TISNO-122 + 123) | mat a his3 leu2 trp1 + REV1p::BenM$^{MP17_H05}$-KI .LEU2 209bp_CYC1p_BenO_ T1::yEGFP-SpHIS5 | This study |
| TISNO-232 | Variant MP04_B01 isolated by FACS (specificity) | FACS isolate (TISNO-122 + 123) | mat a his3 leu2 trp1 + REV1p::BenM$^{MP04_B01}$-KI .LEU2 209bp_CYC1p_BenO_ T1::yEGFP-SpHIS5 | This study |
| TISNO-233 | Variant MP04_B02 isolated by FACS (specificity) | FACS isolate (TISNO-122 + 123) | mat a his3 leu2 trp1 + REV1p::BenM$^{MP04_B02}$-KI .LEU2 209bp_CYC1p_BenO_ T1::yEGFP-SpHIS5 | This study |
| TISNO-234 | Variant MP02_D04 re-transformed (dynamic range) | pTS-105 (MeLS0009) | mat a his3 leu2 trp1 + REV1p::BenM(L104S, T132S, D151E, T157A, I289T)-KI.LEU2 209bp_CYC1p_BenO_ T1::yEGFP-SpHIS5 | This study |
| TISNO-235 | Variant MP02_G10 re-transformed | pTS-106 (MeLS0009) | mat a his3 leu2 trp1 + REV1p::BenM(S279F, Y286N)-KI.LEU2 | This study |

| | | | | |
|---|---|---|---|---|
| | (dynamic range) | | 209bp_CYC1p_BenO_T1::yEGFP-SpHIS5 | |
| TISNO-236 | Variant MP04_B01 re-transformed (specificity) | pTS-107 (MeLS0009) | mat a his3 leu2 trp1 + REV1p::BenM(P201S, T205I, Y293H)-KI.LEU2 209bp_CYC1p_BenO_T1::yEGFP-SpHIS5 | This study |
| TISNO-237 | Variant P2_H02 re-transformed (inversion of function) | pTS-109 (MeLS0009) | mat a his3 leu2 trp1 + REV1p::BenM(L105F, R225K, I240V, T288S, Q291H)-KI.LEU2 209bp_CYC1p_BenO_T1::yEGFP-SpHIS5 | This study |
| TISNO-238 | Variant DAP1_H01 re-transformed (inversion of function) | pTS-110 (MeLS0009) | mat a his3 leu2 trp1 + REV1p::BenM(I108F, K155M, M177R, K220M, L233S, G297D)-KI.LEU2 209bp_CYC1p_BenO_T1::yEGFP-SpHIS5 | This study |
| TISNO-240 | Variant DAP1_H02 re-transformed (inversion of function) | pTS-112 (MeLS0009) | mat a his3 leu2 trp1 + REV1p::BenM(A167T, Q291H)-KI.LEU2 209bp_CYC1p_BenO_T1::yEGFP-SpHIS5 | This study |
| TISNO-241 | Variant MP16_H11 re-transformed (operational range) | pTS-113 (MeLS0009) | mat a his3 leu2 trp1 + REV1p::BenM(S73C, S80N, M126I, E133G, G251C, T299S)-KI.LEU2 209bp_CYC1p_BenO_T1::yEGFP-SpHIS5 | This study |
| TISNO-242 | Variant MP17_F12 re-transformed (operational range) | pTS-114 (MeLS0009) | mat a his3 leu2 trp1 + REV1p::BenM(E226V)-KI.LEU2 209bp_CYC1p_BenO_T1::yEGFP-SpHIS5 | This study |
| TISNO-243 | Variant MP17_H05 re-transformed (operational range) | pTS-116 (MeLS0009) | mat a his3 leu2 trp1 + REV1p::BenM(I154T, M165I, N209K, V258I, T288S, E300K)-KI.LEU2 209bp_CYC1p_BenO_T1::yEGFP-SpHIS5 | This study |
| TISNO-244 | Variant MP17_D08 re-transformed (operational range) | pTS-117 (MeLS0009) | mat a his3 leu2 trp1 + REV1p::BenM(A230V, F253S, Y286N, Y293H)-KI.LEU2 209bp_CYC1p_BenO_T1::yEGFP-SpHIS5 | This study |
| ID7835 | *E. coli* expression strain for variant $BenM^{MP02\_A07}$ | ID7819 (BL21Star(DE3)) | $F^-$*ompT* *hsd*$S_B$ ($r_B$-, $m_B$-) *galdcmrne*131 (DE3) + ID7819 | This study |
| ID7836 | *E. coli* expression strain for variant $BenM^{MP02\_G10}$ | ID7821 (BL21Star(DE3)) | $F^-$*ompT* *hsd*$S_B$ ($r_B$-, $m_B$-) *galdcmrne*131 (DE3) + ID7836 | This study |
| ID7839 | *E. coli* expression strain for variant $BenM^{MP04\_B02}$ | ID7824 (BL21Star(DE3)) | $F^-$*ompT* *hsd*$S_B$ ($r_B$-, $m_B$-) *galdcmrne*131 (DE3) + ID7839 | This study |

| | | | | |
|---|---|---|---|---|
| ID7840 | *E. coli* expression strain for variant $BenM^{MP05\_D07}$ | ID7825 (BL21Star(DE3)) | $F^-$*ompT* *hsd*$S_B$ ($r_B$-, $m_B$-) *galdcmrne*131 (DE3) + ID7840 | This study |
| ID7842 | *E. coli* expression strain for variant $BenM^{DAP1\_H01}$ | ID7827 (BL21Star(DE3)) | $F^-$*ompT* *hsd*$S_B$ ($r_B$-, $m_B$-) *galdcmrne*131 (DE3) + ID7842 | This study |
| ID7845 | *E. coli* expression strain for variant $BenM^{MP16\_H11}$ | ID7830 (BL21Star(DE3)) | $F^-$*ompT* *hsd*$S_B$ ($r_B$-, $m_B$-) *galdcmrne*131 (DE3) + ID7845 | This study |
| ID7846 | *E. coli* expression strain for variant $BenM^{MP17\_F12}$ | ID7831 (BL21Star(DE3)) | $F^-$*ompT* *hsd*$S_B$ ($r_B$-, $m_B$-) *galdcmrne*131 (DE3) + ID7846 | This study |
| ID7849 | *E. coli* expression strain for variant $BenM^{MP17\_D08}$ | ID7834 (BL21Star(DE3)) | $F^-$*ompT* *hsd*$S_B$ ($r_B$-, $m_B$-) *galdcmrne*131 (DE3) + ID7849 | This study |
| JBx_101900 | E. coli biosensing strain for adipic acid | JBx_101898 (DH10B) | $F^-$ *mcrA* *Δ(mrr-hsd*RMS-*mcr*BC*)* *Φ80dlac*ZΔM15 *ΔlacX*74 *end*A1 *rec*A1 *deo*R *Δ(ara,leu)*7697 *ara*D139 *gal*U *gal*K *nup*G *rps*L λ- + JBx_101898 | This study |
| JBx_101901 | E. coli negative control strain for adipic acid biosensing | JBx_101899 (DH10B) | $F^-$ *mcrA* *Δ(mrr-hsd*RMS-*mcr*BC*)* *Φ80dlac*ZΔM15 *ΔlacX*74 *end*A1 *rec*A1 *deo*R *Δ(ara,leu)*7697 *ara*D139 *gal*U *gal*K *nup*G *rps*L λ- + JBx_101899 | This study |

| **Plasmid name** | **Description** | **Source** |
|---|---|---|
| pMeLS0076 | YCp-KlLEU2-REV1p->BenM | Skjoedt et al, 2016 [1] |
| pNic28-Bsa4 | pET expression vector *E. coli* (Addgene plasmid # 26103) | Savitsky et al, 2010 [2] |
| pTS-56 | YCp-KlLEU2-REV1p->BenM(A130D, A153G, P201S, E287V) | This study |
| pTS-66 | YCp-KlLEU2-REV1p->BenM(P201S) | This study |
| pTS-92 | YCp-KlLEU2-REV1p->BenM(A130D) | This study |
| pTS-93 | YCp-KlLEU2-REV1p->BenM(A153G) | This study |
| pTS-94 | YCp-KlLEU2-REV1p->BenM(E287V) | This study |
| pTS-95 | YCp-KlLEU2-REV1p->BenM(P201S, A130D) | This study |
| pTS-96 | YCp-KlLEU2-REV1p->BenM(P201S, A153G) | This study |
| pTS-97 | YCp-KlLEU2-REV1p->BenM(P201S, E287V) | This study |
| pTS-100 | YCp-KlLEU2-REV1p->BenM(H110N, A142T, R160G, T280I) | This study |
| pTS-101 | YCp-KlLEU2-REV1p->BenM(L105M, G138D, K155E, K200E, P201S, N222S, F298S) | This study |

| | | |
|---|---|---|
| pTS-102 | YCp-KlLEU2-REV1p->BenM(E137D, V166I, E277D, Q291H, G297R) | This study |
| pTS-105 | YCp-KlLEU2-REV1p->BenM(L104S, T132S, D151E, T157A, I289T) | This study |
| pTS-106 | YCp-KlLEU2-REV1p->BenM(S279F, Y286N) | This study |
| pTS-107 | YCp-KlLEU2-REV1p->BenM(P201S, T205I, Y293H) | This study |
| pTS-109 | YCp-KlLEU2-REV1p->BenM(L105F, R225K, I240V, T288S, Q291H) | This study |
| pTS-110 | YCp-KlLEU2-REV1p->BenM(I108F, K155M, M177R, K220M, L233S, G297D) | This study |
| pTS-112 | YCp-KlLEU2-REV1p->BenM(A167T, Q291H) | This study |
| pTS-113 | YCp-KlLEU2-REV1p->BenM(S73C, S80N, M126I, E133G, G251C, T299S) | This study |
| pTS-114 | YCp-KlLEU2-REV1p->BenM(E226V) | This study |
| pTS-116 | YCp-KlLEU2-REV1p->BenM(I154T, M165I, N209K, V258I, T288S, E300K) | This study |
| pTS-117 | YCp-KlLEU2-REV1p->BenM(A230V, F253S, Y286N, Y293H) | This study |
| ID7820 | BenM TISNO-124 cloned into pNic28-Bsa4 | This study |
| ID7821 | BenM TISNO-125 cloned into pNic28-Bsa4 | This study |
| ID7824 | BenM-TISNO-128 cloned into pNic28-Bsa4 | This study |
| ID7825 | BenM-TISNO-129 cloned into pNic28-Bsa4 | This study |
| ID7827 | BenM-TISNO-131 cloned into pNic28-Bsa4 | This study |
| ID7830 | BenM-TISNO-134 cloned into pNic28-Bsa4 | This study |
| ID7831 | BenM-TISNO-135 cloned into pNic28-Bsa4 | This study |
| ID7834 | BenM-TISNO-138 cloned into pNic28-Bsa4 | This study |
| JBp_000017 | pBbS5c-RFP | Lee et al., 2011 [3]. |
| JBx_065530 | pSC-gapdhp(EL)-mCherry | Phelan et al., 2017 [4] |
| JBx_101898 | Adipic acid biosensing plasmid | This study |
| JBx_101899 | Negative control biosensing plasmid | This study |

| Primer name | Sequence 5'--> 3' | Description | Source |
|---|---|---|---|
| MelS69-F | GATGAATGCGGCCGCTTTA | Forward primer for random mutagenesis of BenM-EBD | Skjoedt et al, 2016 [1] |
| MelS93-R | CAATACGCCATCAAGTTGCTAAGC | Reverse primer for random mutagenesis of BenM-EBD | Skjoedt et al, 2016 [1] |
| MelS071-F | CTCCTTCCTTTTCGGTTAGAGCGGATGAATGCGGCCGCTTTA | Tailed forward primer for BenM-EBD library assembly by gap repair | Skjoedt et al, 2016 [1] |
| MelS094-R | TCATTTCTTTTACCAATACGCCATCAAGTTGCTAAGC | Tailed reverse primer for BenM-EBD library assembly by gap repair | Skjoedt et al, 2016 [1] |
| TISNO-12F | ACTACGAACTTGCTGATGTCC | Forward BenM sequencing primer (anneals in REV1p) | This study |
| TISNO-13R | CTCAAGCAAGGTTTTCAGTATAATG | Reverse BenM sequencing primer (anneals in CYC1t) | This study |
| ID18723 | GTTGTTCATATGGAGCTGCGCCACCTTCGCTATTTCGTTG | Forward primer for amplifying *E. coli* codon-optimized synthetic DNA constructs BenM (benM-start) | This study |
| ID18724 | GTTGTTGCGGCCGCCCAGTTAGGTGGCTCTGTAAATCCTTC | Reverse primer for amplifying E. coli codon-optimized synthetic DNA constructs BenM (benM-nostop) | This study |
| ID21258 | GTTGTTGCGGCCGCCCAGTTAGGTGGCTCTGTAAAACGTTC | Reverse primer for amplifying E. coli codon-optimized synthetic DNA construct TISNO-129 (benM-rv-tisno129) | This study |
| ID21260 | GTTGTTGCGGCCGCCCAGTTAGGTGGCTCTGTAAAATCTTC | Reverse primer for amplifying E. coli codon-optimized synthetic DNA construct TISNO-131 (benM-rv-tisno131) | This study |
| ID21261 | GTTGTTGCGGCCGCCCAGTTAGGTGGCTCGCTAAATCCTTC | Reverse primer for amplifying E. coli codon-optimized synthetic DNA construct TISNO-134 (benM-rv-tisno134) | This study |

| | | | |
|---|---|---|---|
| ID21262 | GTTGTTGCGGCCGCCCAGTTCTGTGGCTCTGTAAATCCTTC | Reverse primer for amplifying E. coli codon-optimized synthetic DNA construct TISNO-135 (benM-rv-tisno135) | This study |
| j5_00193_(pBbS5c_backbone)_forward | CGAGCTGTACAAGTGAGGATCCAAACTCGAGTAAGGATCTCCAGGC | Forward primer to amplify bacbone adipic acid biosensing plasmid | This study |
| j5_00194_(pBbS5c_backbone)_reverse | CAAATTGGTAAGCGCAACGCAATTAATGTAAGTTAGC | Reverse primer to amplify bacbone adipic acid biosensing plasmid | This study |
| j5_00195_(BenM_Mu_rc)_forward | CATTAATTGCGTTGCGCTTACCAATTTGGTGGTTCAGTAAAACCTTCGTAGGC | Forward primer to amplify BenM$^{TISNO-120}$ from pTS-56 | This study |
| j5_00196_(BenM_Mu_rc)_reverse | ACCTATGGAGTATTTTTAAATGGAATTGAGACACTTGAGATACTTCGTTGCCG | Reverse primer to amplify BenM$^{TISNO-120}$ from pTS-56 | This study |

| **Synthetic DNA name** | **Sequence 5'--> 3'** | **Description** | **Source** |
|---|---|---|---|
| TISNO-113 | /5Biosg/ATACTCCATAGGTATTTTATTATACAAATAATGTGTTTGAACTTATTAAAACATTCTTTTAAGGTATAAACAA | Synthetic 5'end biotinylated (forward) oligo 209bp_*benO*_CYC1 | This study |
| TISNO-110 | TTGTTTATACCTTAAAAGAATGTTTTAATAAGTTCAAACACATTATTTGTATAATAAAATACCTATGGAGTAT | Non-modified (reverse) oligo 209bp_*benO*_CYC1 | This study |
| TISNO-114 | /5Biosg/ATTAGGACCTTTGCAGCATAAATTACTATACTTCTATAGACACACAAACACAAATACACACACTAAATTAATA | Synthetic 5'end biotinylated (forward) oligo 209bp_CYC1 | This study |
| TISNO-112 | TATTAATTTAGTGTGTGTATTTGTGTTTGTGTGTCTATAGAAGTATAGTAATTTATGCTGCAAAGGTCCTAAT | Non-modified (reverse) oligo 209bp_CYC1 | This study |
| TISNO-124 | ATGGAGCTGCGCCACCTTCGCTATTTCGTTGCGGTCGTGGAGGAGCAATCTTTCACCAAGGCTGCCGACAAGTTATGCATCGCACAACCGCCTCTTTCACGTCAAATCCAAAATCTTGAGGAAGAACTGGGCATTCAATTACTTGAACGCGGGTCTCGCCCAGTGAAGACCACCCCCGAGGGGCACTTCTTCTATCAGTATGCCATCAAACTGCTGTCCAACGTTGACCAGATGGTCTCTATGACAAAGCGTATTGCATCTGTAGAGAAGACAATTCGTATTGGCTTTGTTGGCAGCCTGTTGTTTGGCCTGTTACCACGTATTATTAATTTGTACCGTCAGGCGCACCCGAATTTGCGCATTGAACTTTATGAAATGGGTACAAAAGCGCAGACCGAAGCCCTGAAGGAGGGACGCATCGATACTGGATTTGGCCGCTTGAAAATCAGCGATCCAGCAATCAAGCGCACGCTTCTTGGTAACGAGCGCCTGATGGTTGCCGTTCATGCCAGCCACCCACTGAACCAAATGAAGGACAAAGGGGTACACCTTAATGACTTGATTGACGAGAAAATTTTGTTGTATCCTAGTTCTCCGAAGCCAAACTTTTCAACCCACGTAATGAATATCTTCAGTGACCACGGGTTAGAGCCGACCAAAATCAACGAGGTCCGCGAAGTACAGCTTGCTCTGGGGTTGGTGGCAGCCGGAGAGGGGATTAGCCTGGTACCCGCATCTACCCAGTCTATCCAGTTGTTCAATCTGTCATACGTCCCTCTTTTGGACCCTGACGCTATTACCCCCATCTATATCGCCGTGCGTAACATGGAAGAGTCCATTTACATCTATAGCTTATACGAAACAATCCGCCAAATTTATGCTTACGAAGGATTTACAGAGCCACCTAACTGG | BenM$^{MP02_A07}$ codon-optimized for *E. coli* | This study |
| TISNO-125 | ATGGAGCTGCGCCACCTTCGCTATTTCGTTGCGGTCGTGGAGGAGCAATCTTTCACCAAGGCTGCCGACAAGTTATGCATCGCACAACCGCCTCTTTCACGTCAAATCCAAAATCTTGAGGAAGAACTGGGCATTCAATTACTTGAACGCGGGTCTCGCCCAGTGAAGACCACCCCCGAGGGGCACTTCTTCTATCAGTATGCCATCAAACTGCTGTCCAACGTTGACCAGATGGTCTCTATGACAAAGCGTATTGCATCTGTAGAGAAGACAATTCGTATTGGCTTTGTTGGCAGCCTGTTGTTTGGCCTGTTACCACGTATTATTCATTTGTACCGTCAGGCGCACCCGAATTTGCGCATTGAACTTTATGAAATGGGTACAAAAGCGCAGACCGAAGCCCTGAAGGAGGGACGCATCGATGCAGGATTTGGCCGCTTGAAAATCAGCGATCCAGCAATCAAGCGCACGCTTCTTCGTAACGAGCGCCTGATGGTTGCCGTTCATGCCAGCCACCCACTGAACCAAATGAAGGACAAAGGGGTACACCTTAATGACTTGATTGACGAGAAAATTTTGTTGTATCCTAGTTCTCCGAAGCCAAACTTTTCAACCCACGTAATGAATATCTTCAGTGACCACGGGTTAGAGCCGACCAAAATCAACGAGGTCCGCGAAGTACAGCTTGCTCTGGGGTTGGTGGCAGCCGGAGAGGGGATTAGCCTGGTACCCGCATCTACCCAGTCTATCCAGTTGTTCAATCTGTCATACGTCCCTCTTTTGGACCCTGACGCTATT | BenM$^{MP02_G10}$ codon-optimized for *E. coli* | This study |

| | | | |
|---|---|---|---|
| | ACCCCCATCTATATCGCCGTGCGTAACATGGAAGAGTTTACGTACATCTAT<br>AGCTTAAATGAAACAATCCGCCAAATTTATGCTTACGAAGGATTTACAGAG<br>CCACCTAACTGG | | |
| TISNO-128 | ATGGAGCTGCGCCACCTTCGCTATTTCGTTGCGGTCGTGGAGGAGCAAT<br>CTTTCACCAAGGCTGCCGACAAGTTATGCATCGCACAACCGCCTCTTTCA<br>CGTCAAATCCAAAATCTTGAGGAAGAACTGGGCATTCAATTACTTGAACG<br>CGGGTCTCGCCCAGTGAAGACCACCCCCGAGGGGCACTTCTTCTATCAG<br>TATGCCATCAAACTGCTGTCCAACGTTGACCAGATGGTCTCTATGACAAA<br>GCGTATTGCATCTGTAGAGAAGACAATTCGTATTGGCTTTGTTGGCAGCC<br>TGTTGTTTGGCCTGTTACCACGTATTATTCTGTTGTACCGTCAGGCGCACC<br>CGAATTTGCGCATTGAACTTTATGAAATGGGTACAAAAGCGCAGACCGAA<br>GCCCTGAAGGAGGGACGCATCGATGCAGGATTTGGCCGCTTGAAAATCA<br>GCGATCCAGCAATCAAGCGCACGCTTCTTCGTAACGAGCGCCTGATGGTT<br>GCCGTTCATGCCAGCCACCCACTGAACCAAATGAAGGACAAAGGGGTAC<br>ACCTTAATGACTTGATTGACGAGAAAATTTTGTTGTATCCTAGTTCTCCGA<br>AGAGCAACTTTTCAACCCACGTAATGAATATCTTCAGTGACCACGGGTTTG<br>AGCCGACCAAAATCAACGAGGTCCGCGAAGTACAGCTTGCTCTGGGGTT<br>GGTGGCAGCCGGAGAGGGGATTAGCCTGGTACCCGCATCTACCCAGTCT<br>ATCCAGTTGTTCAATCTGTCATACGTCCCTCTTTTGGACCCTGACGCTATT<br>ACCCCCATCTATATCGCCGTGCGTAACATGGAAGAGTCCACGTACATCTA<br>TAGCTTATACGAAACAATCCGCCAAATTTATGCTTACGAAGGATTTACAGA<br>GCCACCTAACTGG | BenM$^{MP04_B02}$ codon-optimized for *E. coli* | This study |
| TISNO-129 | ATGGAGCTGCGCCACCTTCGCTATTTCGTTGCGGTCGTGGAGGAGCAAT<br>CTTTCACCAAGGCTGCCGACAAGTTATGCATCGCACAACCGCCTCTTTCA<br>CGTCAAATCCAAAATCTTGAGGAAGAACTGGGCATTCAATTACTTGAACG<br>CGGGTCTCGCCCAGTGAAGACCACCCCCGAGGGGCACTTCTTCTATCAG<br>TATGCCATCAAACTGCTGTCCAACGTTGACCAGATGGTCTCTATGACAAA<br>GCGTATTGCATCTGTAGAGAAGACAATTCGTATTGGCTTTGTTGGCAGCC<br>TGTTGTTTGGCCTGTTACCACGTATTATTCATTTGTACCGTCAGGCGCACC<br>CGAATTTGCGCATTGAACTTTATGAAATGGGTACAAAAGCGCAGACCGAA<br>GCCCTGAAGGATGGACGCATCGATGCAGGATTTGGCCGCTTGAAAATCA<br>GCGATCCAGCAATCAAGCGCACGCTTCTTCGTAACGAGCGCCTGATGATT<br>GCCGTTCATGCCAGCCACCCACTGAACCAAATGAAGGACAAAGGGGTAC<br>ACCTTAATGACTTGATTGACGAGAAAATTTTGTTGTATCCTAGTTCTCCGA<br>AGCCAAACTTTTCAACCCACGTAATGAATATCTTCAGTGACCACGGGTTAG<br>AGCCGACCAAAATCAACGAGGTCCGCGAAGTACAGCTTGCTCTGGGGTT<br>GGTGGCAGCCGGAGAGGGGATTAGCCTGGTACCCGCATCTACCCAGTCT<br>ATCCAGTTGTTCAATCTGTCATACGTCCCTCTTTTGGACCCTGACGCTATT<br>ACCCCCATCTATATCGCCGTGCGTAACATGGATGAGTCCACGTACATCTA<br>TAGCTTATACGAAACAATCCGCCATATTTATGCTTACGAACGTTTTACAGA<br>GCCACCTAACTGG | BenM$^{MP05_D07}$ codon-optimized for *E. coli* | This study |
| TISNO-131 | ATGGAGCTGCGCCACCTTCGCTATTTCGTTGCGGTCGTGGAGGAGCAAT<br>CTTTCACCAAGGCTGCCGACAAGTTATGCATCGCACAACCGCCTCTTTCA<br>CGTCAAATCCAAAATCTTGAGGAAGAACTGGGCATTCAATTACTTGAACG<br>CGGGTCTCGCCCAGTGAAGACCACCCCCGAGGGGCACTTCTTCTATCAG<br>TATGCCATCAAACTGCTGTCCAACGTTGACCAGATGGTCTCTATGACAAA<br>GCGTATTGCATCTGTAGAGAAGACAATTCGTATTGGCTTTGTTGGCAGCC<br>TGTTGTTTGGCCTGTTACCACGTTTTATTCATTTGTACCGTCAGGCGCACC<br>CGAATTTGCGCATTGAACTTTATGAAATGGGTACAAAAGCGCAGACCGAA<br>GCCCTGAAGGAGGGACGCATCGATGCAGGATTTGGCCGCTTGAAAATCA<br>GCGATCCAGCAATCATGCGCACGCTTCTTCGTAACGAGCGCCTGATGGTT<br>GCCGTTCATGCCAGCCACCCACTGAACCAACGTAAGGACAAAGGGGTAC<br>ACCTTAATGACTTGATTGACGAGAAAATTTTGTTGTATCCTAGTTCTCCGA<br>AGCCAAACTTTTCAACCCACGTAATGAATATCTTCAGTGACCACGGGTTAG<br>AGCCGACCATGATCAACGAGGTCCGCGAAGTACAGCTTGCTCTGGGGAG<br>CGTGGCAGCCGGAGAGGGGATTAGCCTGGTACCCGCATCTACCCAGTCT<br>ATCCAGTTGTTCAATCTGTCATACGTCCCTCTTTTGGACCCTGACGCTATT<br>ACCCCCATCTATATCGCCGTGCGTAACATGGAAGAGTCCACGTACATCTA<br>TAGCTTATACGAAACAATCCGCCAAATTTATGCTTACGAAGATTTTACAGA<br>GCCACCTAACTGG | BenM$^{DAP1_H01}$ codon-optimized for *E. coli* | This study |
| TISNO-134 | ATGGAGCTGCGCCACCTTCGCTATTTCGTTGCGGTCGTGGAGGAGCAAT<br>CTTTCACCAAGGCTGCCGACAAGTTATGCATCGCACAACCGCCTCTTTCA<br>CGTCAAATCCAAAATCTTGAGGAAGAACTGGGCATTCAATTACTTGAACG<br>CGGGTCTCGCCCAGTGAAGACCACCCCCGAGGGGCACTTCTTCTATCAG<br>TATGCCATCAAACTGCTGTGTAACGTTGACCAGATGGTCAATATGACAAA<br>GCGTATTGCATCTGTAGAGAAGACAATTCGTATTGGCTTTGTTGGCAGCC<br>TGTTGTTTGGCCTGTTACCACGTATTATTCATTTGTACCGTCAGGCGCACC<br>CGAATTTGCGCATTGAACTTTATGAAATTGGTACAAAAGCGCAGACCGGT | BenM$^{MP16_H11}$ codon-optimized for *E. coli* | This study |

| | | | |
|---|---|---|---|
| | GCCCTGAAGGAGGGACGCATCGATGCAGGATTTGGCCGCTTGAAAATCA<br>GCGATCCAGCAATCAAGCGCACGCTTCTTCGTAACGAGCGCCTGATGGTT<br>GCCGTTCATGCCAGCCACCCACTGAACCAAATGAAGGACAAAGGGGTAC<br>ACCTTAATGACTTGATTGACGAGAAAATTTTGTTGTATCCTAGTTCTCCGA<br>AGCCAAACTTTTCAACCCACGTAATGAATATCTTCAGTGACCACTGTTTAG<br>AGCCGACCAAAATCAACGAGGTCCGCGAAGTACAGCTTGCTCTGGGGTT<br>GGTGGCAGCCGGAGAGGGGATTAGCCTGGTACCCGCATCTACCCAGTCT<br>ATCCAGTTGTTCAATCTGTCATACGTCCCTCTTTTGGACCCTGACGCTATT<br>ACCCCCATCTATATCGCCGTGCGTAACATGGAAGAGTCCACGTACATCTA<br>TAGCTTATACGAAACAATCCGCCAAATTTATGCTTACGAAGGATTTAGCGA<br>GCCACCTAACTGG | | |
| TISNO-135 | ATGGAGCTGCGCCACCTTCGCTATTTCGTTGCGGTCGTGGAGGAGCAAT<br>CTTTCACCAAGGCTGCCGACAAGTTATGCATCGCACAACCGCCTCTTTCA<br>CGTCAAATCCAAAATCTTGAGGAAGAACTGGGCATTCAATTACTTGAACG<br>CGGGTCTCGCCCAGTGAAGACCACCCCCGAGGGGCACTTCTTCTATCAG<br>TATGCCATCAAACTGCTGTCCAACGTTGACCAGATGGTCTCTATGACAAA<br>GCGTATTGCATCTGTAGAGAAGACAATTCGTATTGGCTTTGTTGGCAGCC<br>TGTTGTTTGGCCTGTTACCACGTATTATTCATTTGTACCGTCAGGCGCACC<br>CGAATTTGCGCATTGAACTTTATGAAATGGGTACAAAAGCGCAGACCGAA<br>GCCCTGAAGGAGGGACGCATCGATGCAGGATTTGGCCGCTTGAAAATCA<br>GCGATCCAGCAATCAAGCGCACGCTTCTTCGTAACGAGCGCCTGATGGTT<br>GCCGTTCATGCCAGCCACCCACTGAACCAAATGAAGGACAAAGGGGTAC<br>ACCTTAATGACTTGATTGACGAGAAAATTTTGTTGTATCCTAGTTCTCCGA<br>AGCCAAACTTTTCAACCCACGTAATGAATATCTTCAGTGACCACGGGTTAG<br>AGCCGACCAAAATCAACGAGGTCCGCGTTGTACAGCTTGCTCTGGGGTT<br>GGTGGCAGCCGGAGAGGGGATTAGCCTGGTACCCGCATCTACCCAGTCT<br>ATCCAGTTGTTCAATCTGTCATACGTCCCTCTTTTGGACCCTGACGCTATT<br>ACCCCCATCTATATCGCCGTGCGTAACATGGAAGAGTCCACGTACATCTA<br>TAGCTTATACGAAACAATCCGCCAAATTTATGCTTACGAAGGATTTACAGA<br>GCCACAGAACTGG | $BenM^{MP17\_F12}$ codon-optimized for *E. coli* | This study |
| TISNO-138 | ATGGAGCTGCGCCACCTTCGCTATTTCGTTGCGGTCGTGGAGGAGCAAT<br>CTTTCACCAAGGCTGCCGACAAGTTATGCATCGCACAACCGCCTCTTTCA<br>CGTCAAATCCAAAATCTTGAGGAAGAACTGGGCATTCAATTACTTGAACG<br>CGGGTCTCGCCCAGTGAAGACCACCCCCGAGGGGCACTTCTTCTATCAG<br>TATGCCATCAAACTGCTGTCCAACGTTGACCAGATGGTCTCTATGACAAA<br>GCGTATTGCATCTGTAGAGAAGACAATTCGTATTGGCTTTGTTGGCAGCC<br>TGTTGTTTGGCCTGTTACCACGTATTATTCATTTGTACCGTCAGGCGCACC<br>CGAATTTGCGCATTGAACTTTATGAAATGGGTACAAAAGCGCAGACCGAA<br>GCCCTGAAGGAGGGACGCATCGATGCAGGATTTGGCCGCTTGAAAATCA<br>GCGATCCAGCAATCAAGCGCACGCTTCTTCGTAACGAGCGCCTGATGGTT<br>GCCGTTCATGCCAGCCACCCACTGAACCAAATGAAGGACAAAGGGGTAC<br>ACCTTAATGACTTGATTGACGAGAAAATTTTGTTGTATCCTAGTTCTCCGA<br>AGCCAAACTTTTCAACCCACGTAATGAATATCTTCAGTGACCACGGGTTAG<br>AGCCGACCAAAATCAACGAGGTCCGCGAAGTACAGCTTGTTCTGGGGTT<br>GGTGGCAGCCGGAGAGGGGATTAGCCTGGTACCCGCATCTACCCAGTCT<br>ATCCAGTTGAGCAATCTGTCATACGTCCCTCTTTTGGACCCTGACGCTATT<br>ACCCCCATCTATATCGCCGTGCGTAACATGGAAGAGTCCACGTACATCTA<br>TAGCTTAAATGAAACAATCCGCCAAATTCATGCTTACGAAGGATTTACAGA<br>GCCACCTAACTGG | $BenM^{MP17\_D08}$ codon-optimized for *E. coli* | This study |
| BP-01 | TCTCAAGTGTCTCAATTCCATTTAAAAATACTCCATAGGTATTTTATTATAC<br>AAATAATGTGTTTGAACTTATTAAAACATTCTTTTAAGGTATAAACAAGCAA<br>GAAAGACAAGAAGAAGGCAGGGGCTTGACCCATTAAATGCTTTCTTCAAT<br>TTGGAAAATTGAAAGCTGAAATGGATATTCGTTTTATTTGTCGGTTCTGCC<br>GTTAAGTAAACATTTTATGCGTTGCGTTGTTTAATTGAATGTTTGACTAAGC<br>ACAGCGTTTTGCTCTGGCCTAGACAAGTTTCTTATTTTGGAATGTTGGAGA<br>AAGGATATGGTCTCCAAGGGCG | 0.3 kb DNA binding site of BenM (benO) with overlap regions | This study |
